# Detection of Rapidly Accumulating Stress-Induced SUMO in Prostate Cancer Cells by a Fluorescent SUMO Biosensor

**DOI:** 10.1101/2021.01.22.427833

**Authors:** Rui Yin, Jiacheng Song, Aurora Esquela-Kerscher, Oliver Kerscher

## Abstract

SUMO conjugates and SUMO chains form when SUMO, a small ubiquitin-like modifier protein, is covalently linked to other cellular proteins or itself. During unperturbed growth, cells maintain balanced levels of SUMO conjugates. In contrast, eukaryotic cells that are exposed to proteotoxic and genotoxic insults mount a cytoprotective SUMO-Stress Response (SSR). One hallmark of the SSR is a rapid and massive increase of SUMO conjugates in response to oxidative, thermal, and osmotic stress. Here, we use a recombinant fluorescent SUMO biosensor, KmUTAG-fl, to investigate differences in the SSR in a normal human prostate epithelial cell line immortalized with SV40 (PNT2) and two human prostate cancer cell lines that differ in aggressiveness and response to androgen (LNCaP and PC3). In cells that grow unperturbed, SUMO is enriched in the nuclei of all three cell lines. However, upon 30 minutes of exposure to ultraviolet radiation (UV) or oxidative stress, we detected significant cytosolic enrichment of SUMO as measured by KmUTAG-fl staining. This rapid enrichment in cytosolic SUMO levels was on average 5-fold higher in the LNCaP and PC3 prostate cancer cell lines compared to normal immortalized PNT2 cells. Additionally, this enhanced enrichment of cytosolic SUMO was reversible as cells recovered from stress exposure. Our study validates the use of the fluorescent KmUTAG-fl SUMO biosensor to detect differences of SUMO levels and localization between normal and cancer cells and provides new evidence that cancer cells may exhibit an enhanced SSR.

## Introduction

Sumoylation, the protein modification with the small ubiquitin-like modifier (SUMO), is a fundamental protein modification process that is conserved from yeast to humans and controls the function, activation, localization, interaction, and half-life of specific cellular proteins (reviewed by Z. Zhao et al., 2018). The proteins that promote sumoylation and SUMO dynamics act via a cascade of SUMO activating (E1), conjugating (E2), and ligase (E3) enzymes as well as SUMO proteases and SUMO-targeted ubiquitin ligases (Kerscher, Felberbaum, & Hochstrasser, 2006). A recent study identified more than 4,300 sumoylation sites in more than 1,600 proteins from HeLa cells. This study confirmed that sumoylation is a predominantly nuclear event that regulates transcription, DNA repair, chromatin remodeling, pre-mRNA splicing and ribosome assembly. Of note, a large number of human disease proteins were found to be targets of SUMO modification (Hendriks et al., 2014; Sarge & Park-Sarge, 2009).

An early study by Saitoh and Hinchey showed that exposure to proteotoxic and genotoxic stressors led to an extremely rapid increase of SUMO-conjugated proteins in the cell, often within minutes (Saitoh & Hinchey, 2000). For example, SUMO-modification is rapidly enhanced when cells are subjected to reagents and conditions that damage proteins (e.g. hydrogen peroxide and increased temperature) or DNA (e.g. UV irradiation). This phenomenon is now termed the SUMO stress response or SSR (Lewicki, Srikumar, Johnson, & Raught, 2015). A long list of sumoylated proteins that accumulate in response to stress, including many chromatin remodeling factors and transcription factors, has been identified using proteomic approaches (reviewed in (Golebiowski et al., 2009 and references therein). It remains unclear however, why this massive SUMO modification event unfolds as cells experience stress.

There is good evidence that organisms as diverse as yeast and humans utilize SUMO modification in their cellular stress response. This is borne out by the reduced stress tolerance of cells that lack intact sumoylation pathways. In yeast, a number of non-essential genes involved in the response to proteotoxic and genotoxic stress result in lethality when paired with genetic defects in sumoylation and desumoylation (reviewed in Seeler & Dejean, 2017). Correspondingly, mammalian cells that are depleted of SUMO or the SUMO protease SENP1 show reduced ability to survive acute heat shock or exposure to ionizing radiation (Golebiowski et al., 2009; R.-T. Wang, Zhi, Zhang, & Zhang, 2013b). This suggests that sumoylation plays an important role in the response to proteotoxic and genotoxic stress.

SUMO’s role in stress tolerance, however, remains enigmatic at best. Findings from three recent studies arrived at different, yet non-mutually exclusive conclusions (Golebiowski et al., 2009; Lewicki et al., 2015; Liebelt et al., 2019). One study found that, in mammalian cells, SUMO isoforms served a chaperone-like function and maintained the homeostasis of large chromatin-associated nuclear proteins during stress (Seifert, Schofield, Barton, & Hay, 2015). Another found that sumoylation temporarily stabilized denatured proteins after heat shock, preventing them from aggregating before proteasome-mediated degradation (Liebelt et al., 2019). A third study found that environmental stress induced a wave of transcription-coupled sumoylation and the SSR was found to be blocked when transcription was inhibited (Lewicki et al., 2015).

A rapid increase in sumoylation and the SSR occurs due to increased activity of SUMO E3 ligases, possibly coupled with a decrease in SUMO protease activity. This is borne out by the finding that initiation of the SSR in stressed cells is caused primarily by the E3 SUMO ligase Siz1 in yeast, and the combined effort of the orthologous SUMO ligases PIAS1-4 in heat-stressed osteosarcoma cells (Lewicki et al., 2015; Seifert et al., 2015). Additionally, several SUMO proteases (Ulp1 in yeast and SENP 1,2,3,7 in mammalian cells) are inactivated by heat and/or oxidative stress, suggesting they may act as stress sensors (Pinto et al., 2012). Inactivation of these SUMO proteases invariably results in an accumulation of SUMO-conjugated proteins as well as SUMO chains. Recent research from our lab indicates that the *S. cerevisiae* SUMO protease Ulp1 is unable to bind SUMO in the presence of extremely low levels of hydrogen peroxide [1.8mM], underscoring a potential role of SUMO proteases as redox sensors (Peek et al., 2018). One notable exception, the mammalian SUMO protease SENP6 is not inactivated by heat stress and becomes recruited to chromatin (Pinto et al., 2012; Seifert et al., 2015). SENP6 is similar to the yeast SUMO protease Ulp2, which in turn is required for recovery from the SSR in yeast. This suggests that both SENP6 and ULP2 are involved in removing SUMO chains as a necessary step for the recovery of cells from the SSR (Lewicki et al., 2015). Additionally, low levels of oxidative stress also rapidly disable the SUMO E1 (Uba2) and E2 (Ubc9) enzymes – via the formation of a disulfide bond between their catalytic cysteine residues – raising the question of how sumoylation can become amplified dramatically during the SSR when SUMO conjugation is halted (Bossis & Melchior, 2006). The answer is unknown, but a pre-existing pool of Ubc9 with non-covalently bound SUMO, which has been shown to be involved in the formation of SUMO chains, may hint at the answer (Knipscheer, van Dijk, Olsen, Mann, & Sixma, 2007).

The SSR pathway exists in single cell eukaryotes (e.g. yeasts), in normal mammalian cells, and is generally dysregulated in cancer cells (Seeler & Dejean, 2017). Cancer cells are subjected to a host of adverse conditions including hypoxic environments within tumors, attack by tumor-invading immune cells, and rampant aneuploidies that dysregulate cellular proteostasis. Consequently, cancer cells rely on enhanced stress response pathways to survive under these conditions. In general, sumoylation enzymes, such as activating (E1) and conjugating (E2) enzymes have been found to be elevated in tumors, potentially altering the activity of dozens of SUMO-modified tumor suppressors, oncoproteins, and stress response proteins, including heat-shock proteins (e.g. Hsp90), hypoxia-inducible factors (e.g Hif1A), and inflammatory signaling factors (e.g IkBalpha) (reviewed in (Seeler & Dejean, 2017)). Specifically, the overexpression of the SUMO ligase Ubc9 in cancers has been linked to poor treatment outcomes and Ubc9 overexpression may be a useful biomarker for cervical cancer (Mattoscio, 2015; Wu, Zhu, Ding, Beck, & Mo, 2009). However, elevated levels of some SUMO proteases have also been linked to breast and prostate cancer development (Karami et al., 2017; Q. Wang et al., 2013a). These examples suggest that accelerated SUMO dynamics, both sumoylation and desumoylation, are at play in the stress resilience of cancer cells.

We recently developed a novel SUMO-trapping protein from a thermotolerant yeast, *Kluyveromyces marxianus (Km)* (Peek et al., 2018). Specifically, we generated a catalytically inactive recombinant SUMO-trapping fragment of the UD domain of Ulp1 (the yeast SUMO protease), termed KmUTAG. KmUTAG can efficiently bind to a variety of purified SUMO isoforms and immobilized SUMO1 with nanomolar affinity (Kd∼12.8nM). Moreover, KmUTAG has high affinity for SUMO and SUMO-modified proteins even in the presence of oxidative stressors (170mM H_2_O_2_), reducing agents (5mM TCEP Tris(2-carboxyethyl)phosphine hydrochloride), denaturants (up to 2M UREA), and temperature stress (42°C) that induce protein misfolding (Peek et al., 2018).

To visualize SUMO within cells with the KmUTAG, we developed a Recombinant Fluorescent SUMO biosensor KmUTAG-fl (Yin, Harvey, Shakes, & Kerscher, 2019). KmUTAG-fl is a recombinant mCherry-tagged SUMO-trapping fusion protein. This stress-tolerant pan-SUMO specific biosensor is produced recombinantly in bacteria. Once purified it recognizes and traps native SUMO-conjugated proteins and SUMO chains in fixed permeablized cells. This biosensor protein compares favorably to staining protocols with SUMO specific antibodies as it has reduced affinity for free, unconjugated SUMO. Meanwhile, it can be used to analyze SUMO variants from additional model and non-model systems. In the present study, we used KmUTAG-fl to detect SUMO in a variety of human prostate cells lines; a non-tumorigenic SV40-immortalized prostate epithelial cell line (PNT2), an aggressive androgen-insensitive prostate cancer cell line derived from bone metastasis with high metastatic potential (PC3), and a hormone-sensitive metastatic prostate cancer cell line derived from lymph node metastasis with low metastatic potential (LNCaP). Using the fluorescent KmUTAG-fl biosensor, we compared SUMO levels in the nuclei and extra-nuclear compartment (cytosol) in untreated, UV-treated, and H_2_0_2_-treated cells. We detected significant differences in the SUMO profiles between normal (from here on defined as non-malignant, immortalized PNT2 prostate epithelial cells) and cancer prostate cancer cells. After stress exposure, both prostate cancer cell lines showed a cytosolic SUMO enrichment that was ∼5-fold higher than normal PNT2 cells. The cytosolic SUMO enrichment was detected within 30 min of stress exposure and was completely reversible after recovery in fresh media. While there was a clear difference between cancer and normal prostate cells, we did not detect a difference between cells exhibiting low (LNCaP) and high (PC3) metastatic potential. Our limited study suggests that differences in the SSR may be linked to the enhanced robustness of cancer cells and therefore, SUMO profile visualization using the KmUTAG-fl biosensor could be a useful tool to differentiate normal and tumorigenic cells.

## RESULTS

### Cytosolic SUMO levels increased in PC3 prostate cancer cells upon UV irradiation

We previously showed that KmUTAG-fl is a single-chain recombinant pan-SUMO binding protein that reliably detected SUMO conjugates in fixed nematode gonads and in mammalian cells (Yin et al., 2019). In unperturbed mammalian cells, the KmUTAG-fl signal co-localized with SUMO2 in the nucleoplasm and in distinct nuclear foci (Peek et al., 2018; Yin et al., 2019). To investigate the differences of the SSR in normal and cancer cells, we used KmUTAG-fl to investigate the impact of UV-induced stress on SUMO levels in two human prostate cell lines, immortalized normal PNT2 cells derived from prostate epithelium and PC3 adenocarcinoma cells with high metastatic potential (Berthon, Cussenot, Hopwood, Leduc, & Maitland, 1995; Tai et al., 2011). PC3 and PNT2 cells were grown on coverslips and subjected to various doses of UV irradiation (50 mJ/m^2^ and 150 mJ/m^2^). After irradiation, the cells were allowed to recover for 30 min in culture medium before fixing and staining with KmUTAG-fl. A rapid increase in KmUTAG-fl staining after UV exposure (150 mJ/m^2^) was visually apparent in PC3 cells (Fig. 1A left panel), but was not discernible in PNT2 cells (Fig. 1A right panel). Levels of KmUTAG-fl in PC3 cells were measured and quantified using CellProfiler, revealing a significant increase in SUMO levels between the un-irradiated (zero “0” mJ/m^2^) and irradiated (150 mJ/m^2^) samples (Fig. 1B top panels). Specifically, we found that after UV treatment (150 mJ/m2) KmUTAG-fl staining in the cytosol (denoting the extra-nuclear region of the cell) increased by ∼54% in PC3 cells (Fig. 1C). In comparison, no significant change in SUMO accumulation was detected in the cytosol of PNT2 (Fig. 1B bottom left panel). Therefore, the difference in cytosolic SUMO levels between UV-treated PC3 and PNT2 cells (150 mJ/m2) was approximately 14-fold. Concomitantly, UV-treatment also increased SUMO levels in the nucleus of PC3 cells, albeit only by 27% (Fig. 1B top right panel). No significant change in SUMO accumulation was detected in nuclei of PNT2 (Fig. 1B bottom right panel). Importantly, taking into account nuclear and cytosolic KmUTAG-fl signals, we recorded an increasing relative cytosolic enrichment (RCE) of SUMO after UV irradiation (Fig. 1D). These data indicated that the SSR in PC3 cancer cells involved a significant increase of SUMO or SUMO conjugates in the cytosol.

**Figure 1.**
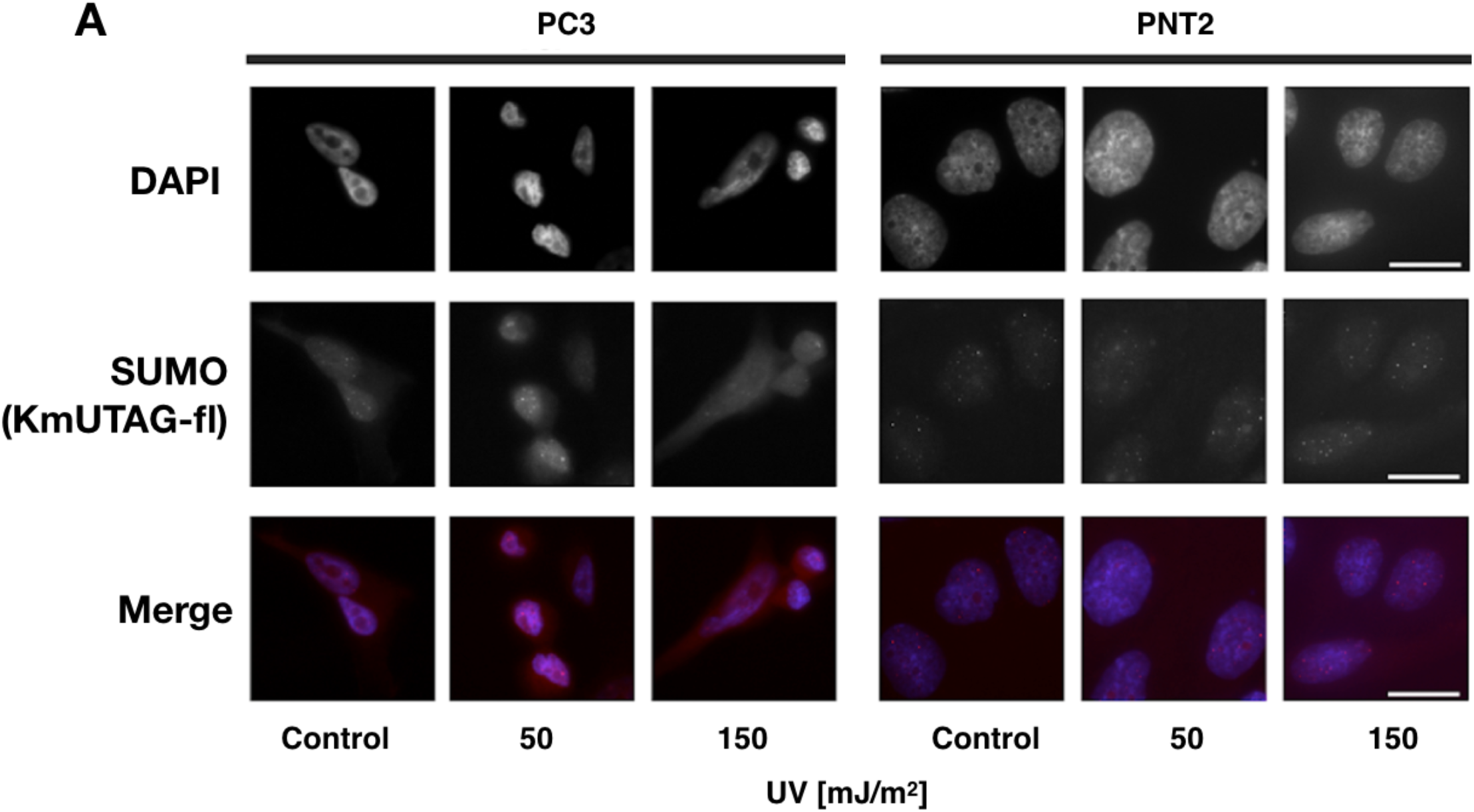
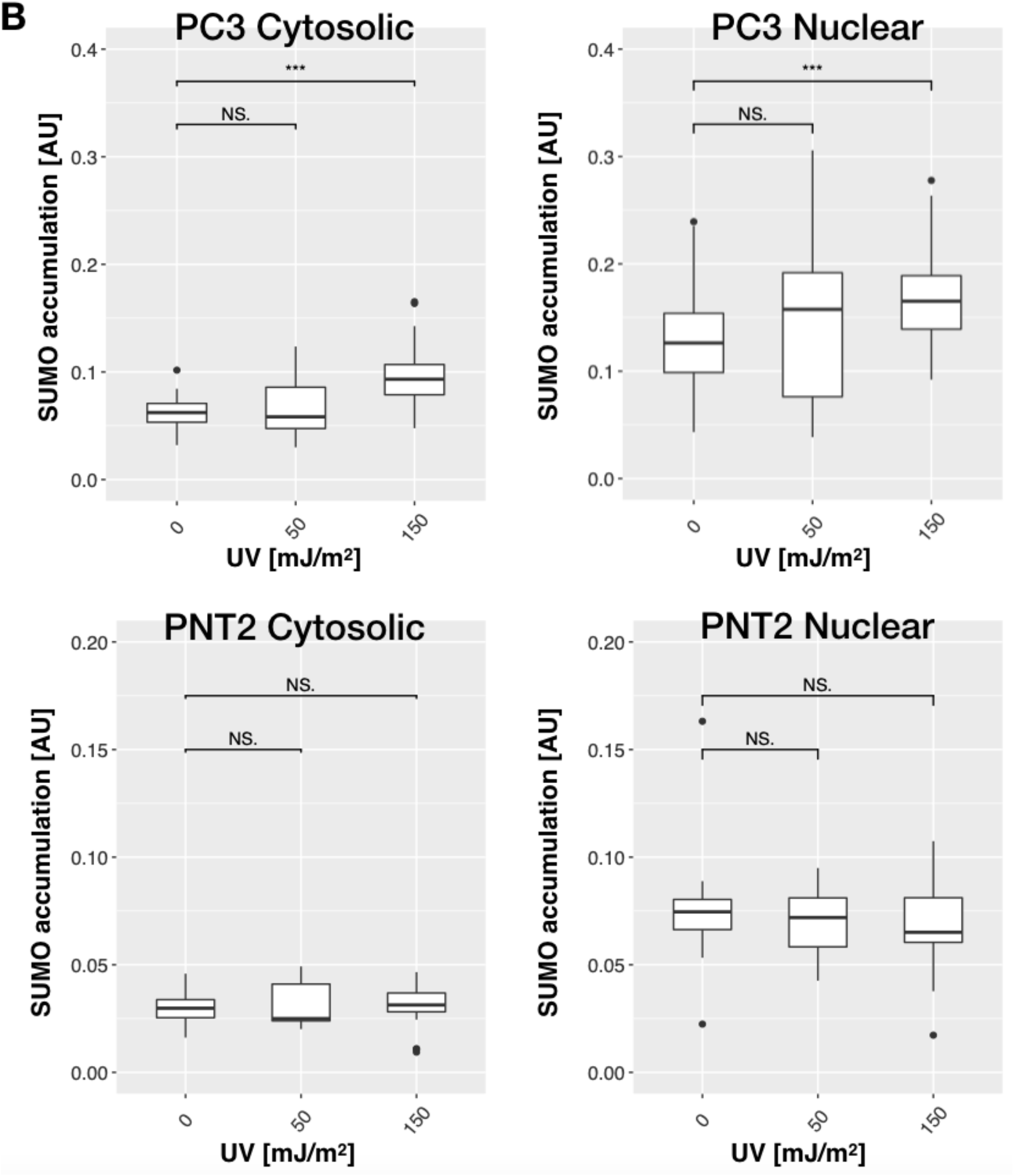
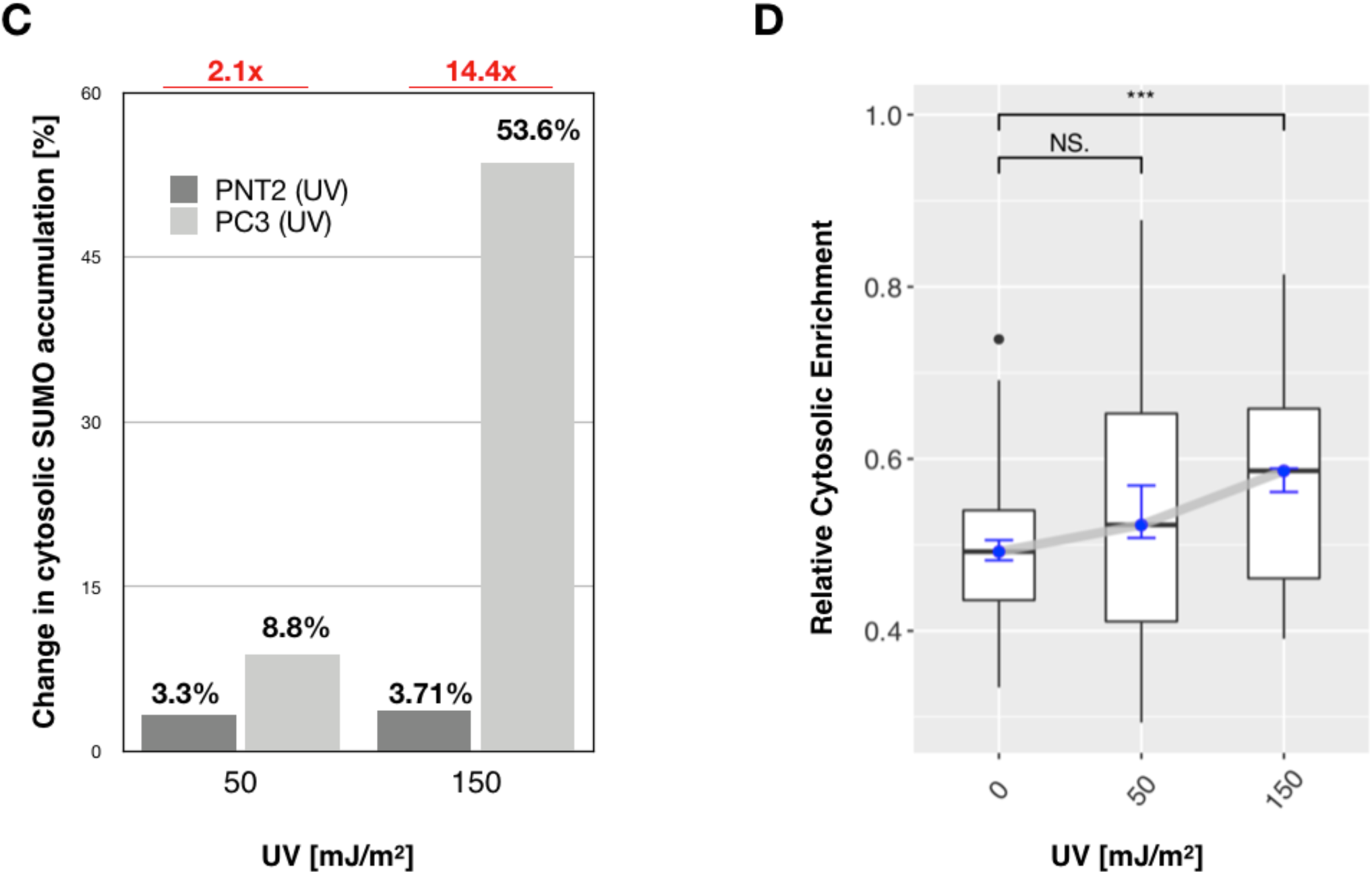
SUMO enrichment of UV irradiated PC3 and PNT2 cells. **(A)** UV dosage changed the SUMO profile of PC3 cells but not of PNT2 cells. UV irradiated PC3 and PNT2 cells (0 (Control), 50 mJ/m^2^, and 150 mJ/m^2^) were stained with KmUTAG-fl (SUMO), and visualized using confocal microscopy for mCherry (KmUTAG-fl, red) and DAPI (blue). **(B)** Nuclear and cytosolic KmUTAG-fl signal intensities in PC3 and PNT2 cells were quantified with CellProfiler [Carpenter et al., 2006 PMID: 17076895]. PC3: 0 mJ/m^2^, n=54; 50 mJ/m^2^, n=27; 150 mJ/m^2^, n=68. PNT2: 0 mJ/m^2^, n=20; 50 mJ/m^2^, n=20; P 150 mJ/m^2^, n=29. **(C)** Increased cytosolic SUMO accumulation (KmUTAG-fl signal intensity) of PC3 cells compared to PNT2 cells. Intensity change was calculated as the difference between the UV irradiated and untreated Control cells. **(D)** Relative Cytosolic Enrichment (RCE) ratio of SUMO (KmUTAG-fl signal) in PC3 cells increased with UV dosage. The RCE was calculated as the ratio between cytosolic and nuclear fluorescence intensity. Statistical analysis by unpaired t-test (NS: P > 0.05, * P ≤ 0.05, ** P ≤ 0.01, *** P ≤ 0.001). Scale bars: 20 µm.

Next, we wanted to know whether this phenomenon of increased extra-nuclear SUMO accumulation could be recapitulated using the monoclonal anti-SUMO2 8A2 antibody to visualize SUMO (X.-D. Zhang et al., 2008). Therefore, we compared the ability of KmUTAG-fl and the anti-SUMO2 8A2 antibody to detect a change in the localization and levels of SUMO following UV-irradiation of PC3 cells. As expected, non-irradiated PC3 cells stained with the 8A2 antibody revealed SUMO2 localization in the nucleus. However, upon UV irradiation [250mJ/m^2^], SUMO2 was also detected in the cytosol of these cells (Fig. 2A). Quantitation of KmUTAG-fl and 8A2 signal revealed that cytosolic SUMO/SUMO2 levels increased by 67% and 52%, respectively (Fig. 2C). However, unlike the KmUTAG-fl stained sample, 8A2 antibody staining did not reveal a significant change of nuclear SUMO levels in irradiated PC3 cells (Fig. 2B left panel). In summary, these data confirmed that KmUTAG-fl was an effective reagent to detect increased SUMO levels, especially in the cytosol of PC3 cancer cells after acute UV exposure.

**Figure 2:**
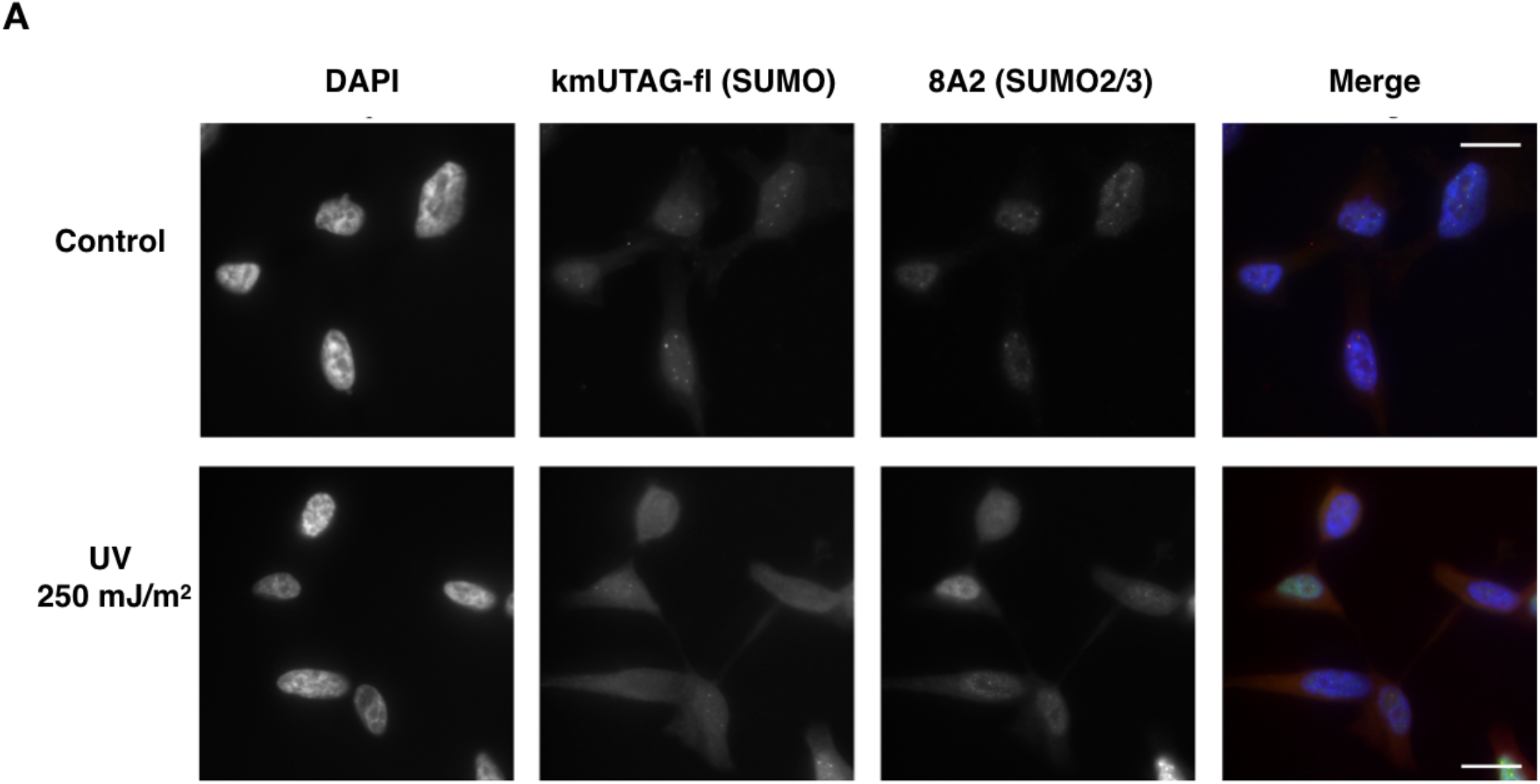
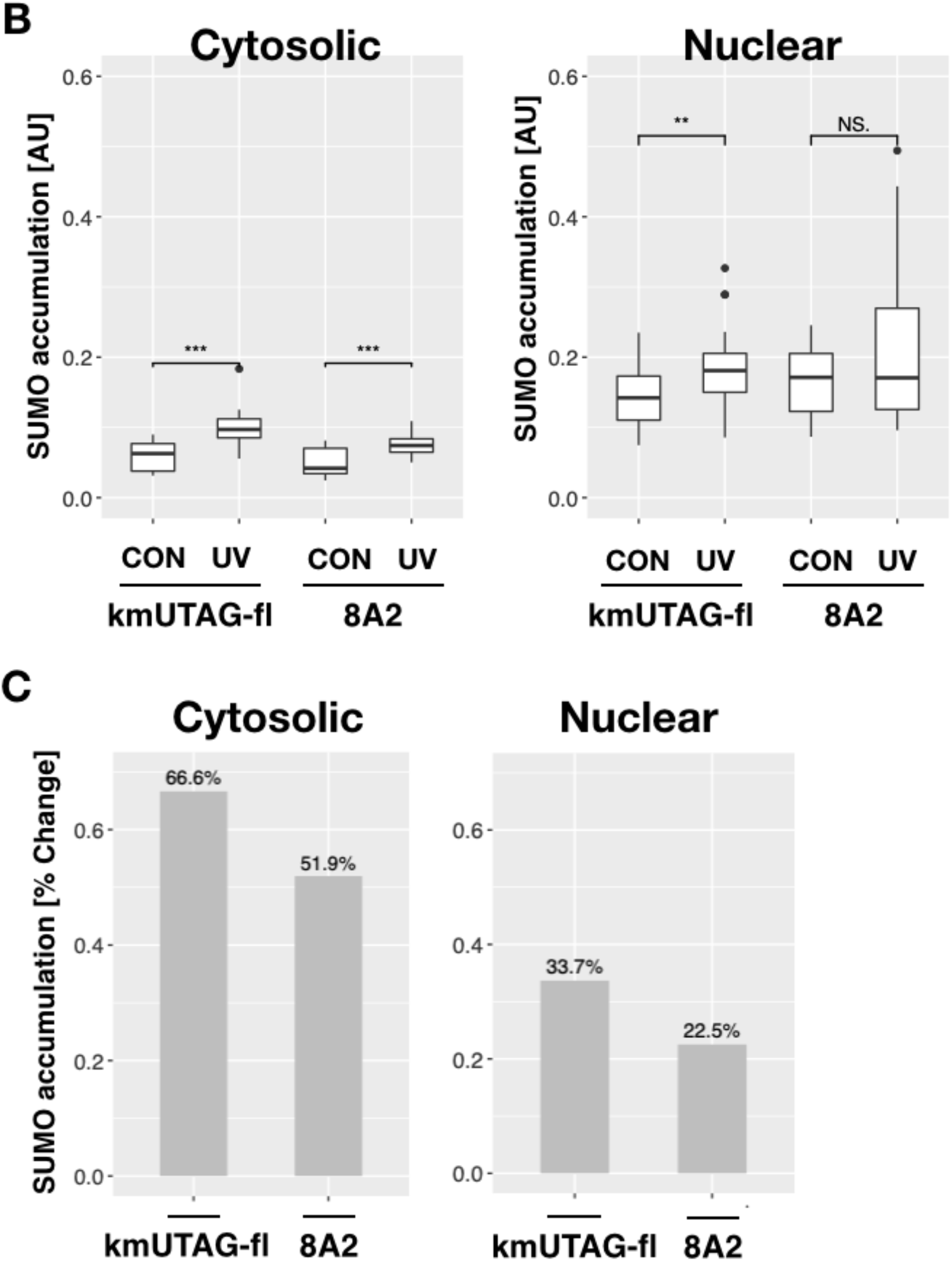
KmUTAG-fl and anti-SUMO2 8A2 antibody co-staining of UV irradiated cancer cells. **(A)** Similar SUMO profiles were observed using the KmUTAG-fl biosensor (red) and the anti-SUMO antibody (green). PC3 cells were irradiated with 250 mJ/m^2^ UV. Fixed cells were co-stained with KmUTAG-fl (SUMO), anti-SUMO2 8A2 antibody (SUMO2), and DAPI (nucleus, blue). Control cells were not UV-irradiated. Scale bars = 20 µm. **(B)** Average cytosolic and nuclear KmUTAG-fl and 8A2 signal intensities (SUMO accumulation) were quantified with CellProfiler (km_Control n=23; km_UV n=22; antiS2_Control n=23; antiS2_UV n=22). AU: arbitrary units. **(C)** Percent change in cytosolic and nuclear KmUTAG-fl and antibody signal intensity in PC3 cells. Statistical analysis by unpaired t-test (NS: P > 0.05, * P ≤ 0.05, ** P ≤ 0.01, *** P ≤ 0.001). Scale bars: 20 µm.

### Cytosolic SUMO levels increased in PC3 prostate cancer cells due to oxidative stress

The rapid accumulation of extra-nuclear SUMO levels due to UV irradiation prompted us to investigate differences in the SSR between PC3 and PNT2 cells after exposure to oxidative stress. PC3 and PNT2 cells were treated with varying concentrations of hydrogen peroxide (0.5µM to 30mM H_2_O_2_) for 30 min, and immediately fixed and stained with KmUTAG-fl. Our analysis of PC3 cells revealed a significant increase of cytosolic and nuclear KmUTAG-fl signal after treatment with 25µM to 30mM H_2_O_2_ (Fig. 3A and B). The most significant median increase of SUMO staining intensity in PC3 cells was observed after 1mM H_2_O_2_ treatment both in the cytosol (216%) and nucleus (87%) (Fig. 3B and 3E). By comparison, the cytosolic and nuclear KmUTAG-fl signal in PNT2 cells did not reveal a steady trend of increasing SUMO signal following treatment (Fig. 3C and D). Rather, a statistically significant increase in cytosolic PNT2 SUMO staining intensity after treatment with 20µM H_2_O_2_ (∼40%) was observed while cytosolic and nuclear SUMO accumulation fell below the intensity of the untreated control (28% reduction) at 5mM H_2_O_2_ (Fig. 3D and E). Taking into account trends of both nuclear and cytosolic KmUTAG-fl signals, we find an increasing RCE of SUMO after H_2_O_2_ treatment that is specific to PC3 cells (Fig. 3F and G).

**Figure 3:**
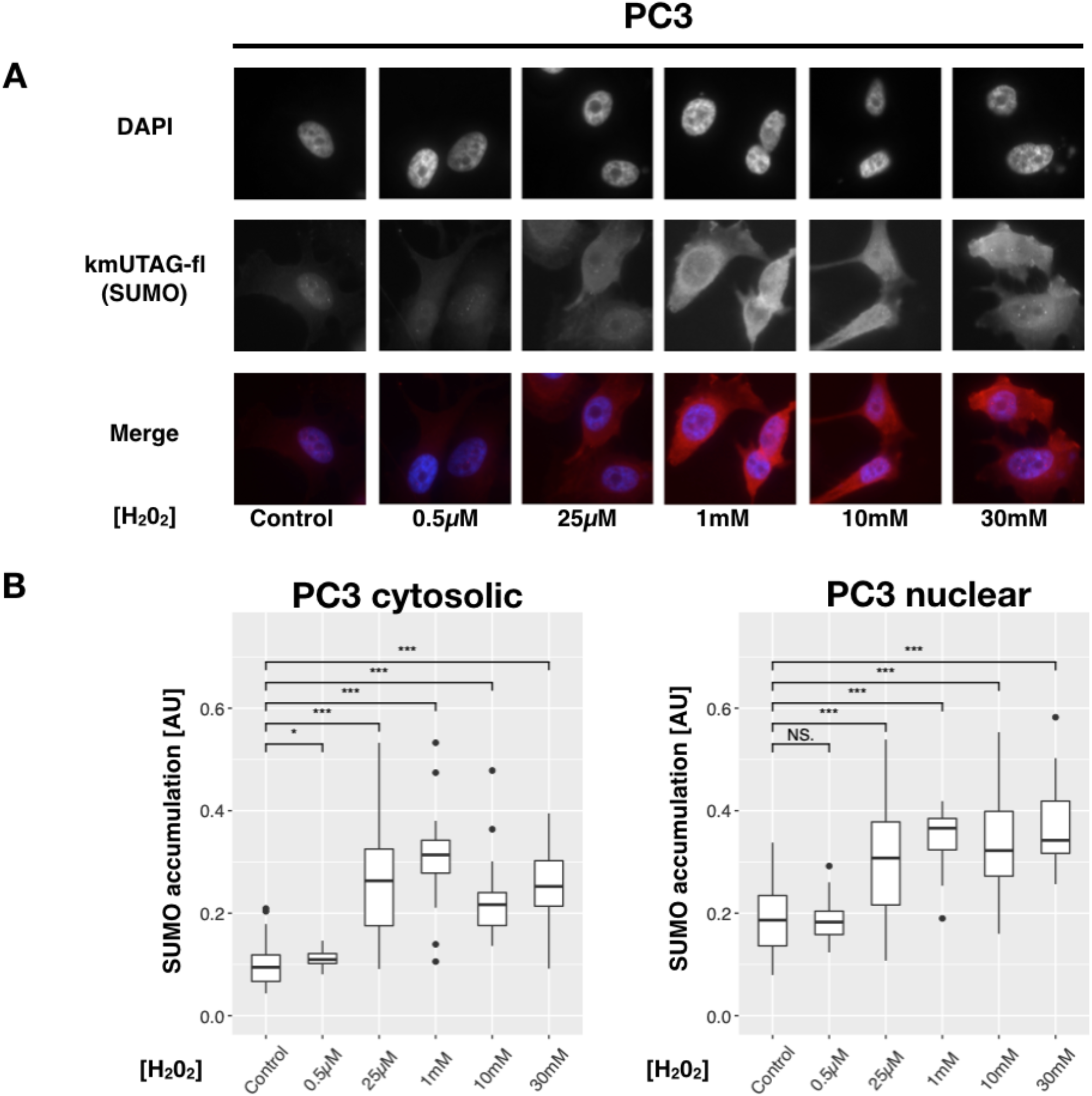
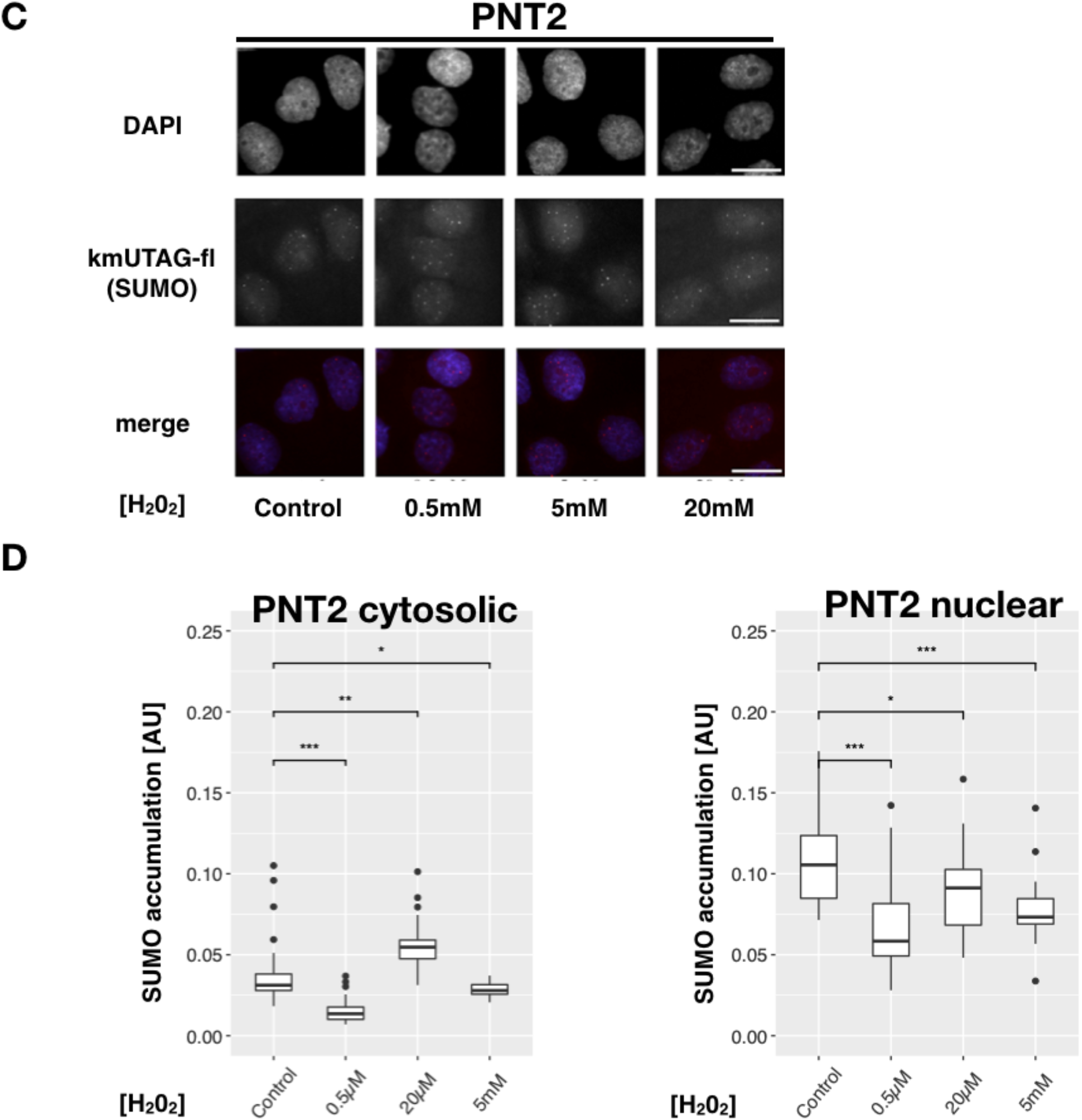
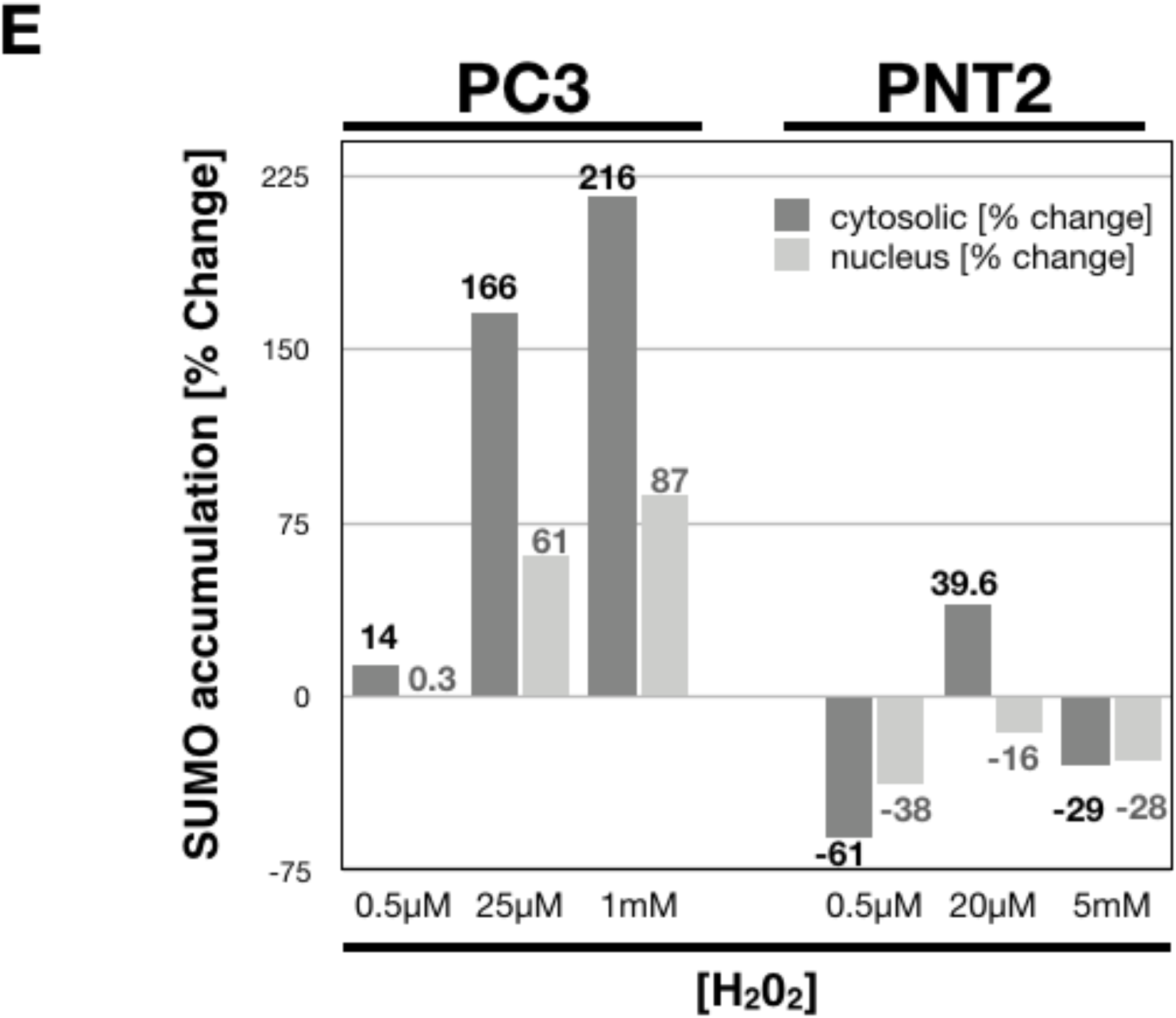
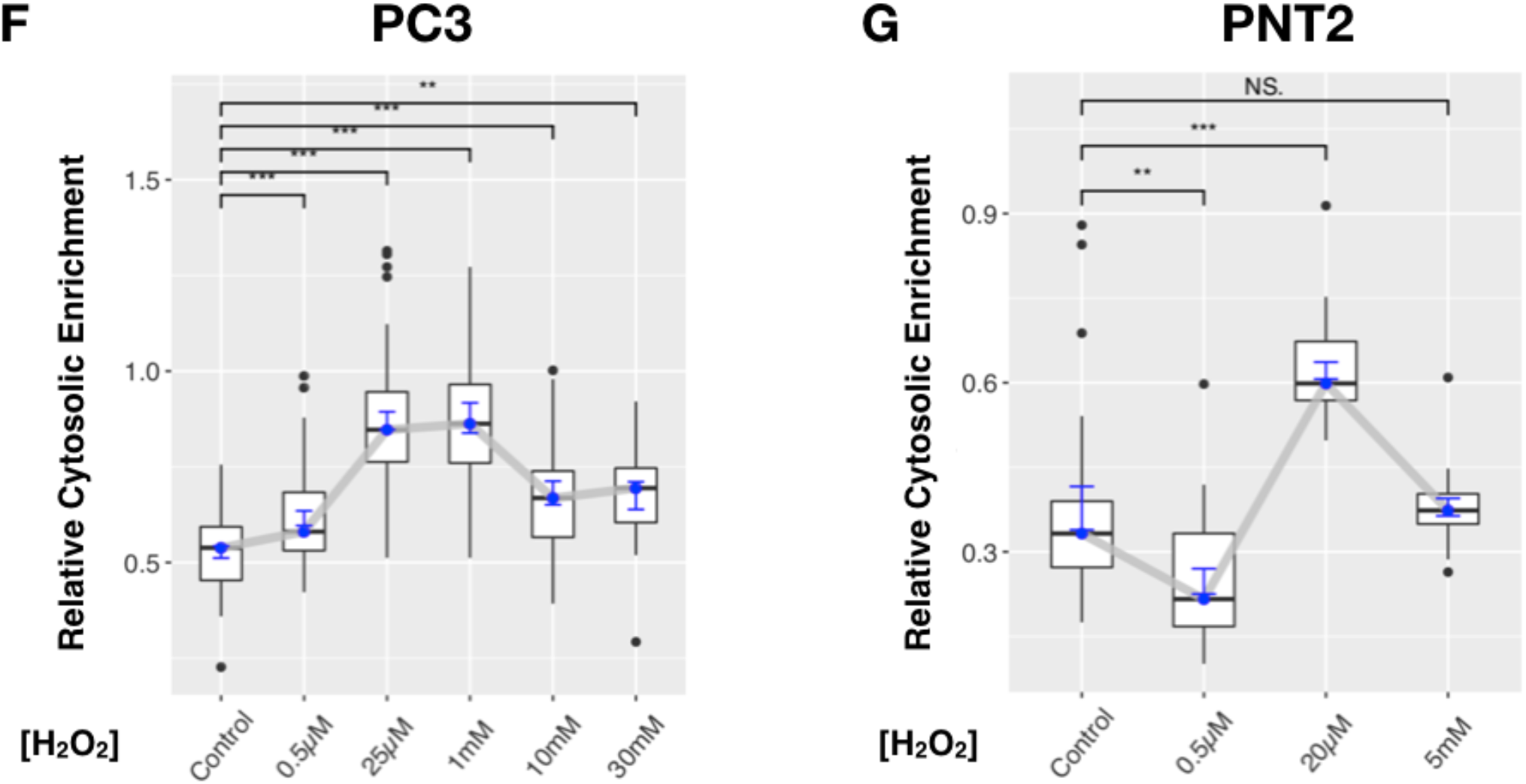
SUMO enrichment of H_2_O_2_ treated PC3 and PNT2 cells. **(A)** H_2_O_2_ treated (0.5µM, 25µM, 1mM, 10mM, 30mM) or control PC3 cells were fixed and stained for SUMO with KmUTAG-fl, visualized and quantified as previously described. **(B)** Average cytosolic and nuclear KmUTAG-fl signal intensity (SUMO accumulation) were quantified with CellProfiler (PC3 Control n=47; PC3 0.5µM n=44, PC3 25µM n=54; PC3 1mM n=23; PC3 10mM n=25; PC3 30mM n=17) **(C)** H_2_O_2_ treated (0.5mM, 5mM, 20mM) or control PNT2 cells were fixed and stained for SUMO with KmUTAG-fl, visualized and quantified as above. **(D)** Average cytosolic and nuclear KmUTAG-fl (SUMO accumulation) were quantified with CellProfiler PNT2 Control n=24; PNT2 0.5µM n=27, PNT2 20µM n=33; PNT2 5mM n=20). **(E)** Increased cytosolic and nuclear SUMO accumulation (KmUTAG-fl signal intensity) of PC3 cells compared to PNT2 cells. Intensity change was calculated as the difference between the H_2_O_2_ treated and untreated Control cells. **(F)** Relative Cytosolic Enrichment (RCE) ratio of SUMO (KmUTAG-fl signal) in PC3 cells increased with H_2_O_2_ concentration. The RCE was calculated as the ratio between cytosolic and nuclear fluorescence intensity. **(G)** Variability in the Relative Cytosolic Enrichment (RCE) ratio of SUMO (KmUTAG-fl signal) in PNT2 cells at various H_2_O_2_ concentrations. The RCE was calculated as the ratio between cytosolic and nuclear fluorescence intensity. Statistical analysis by unpaired t-test (NS: P > 0.05, * P ≤ 0.05, ** P ≤ 0.01, *** P ≤ 0.001). Scale bars: 20 µm.

Simultaneously, we also quantitated nuclear SUMO foci in H_2_O_2_-treated PNT2 and PC3 cells. Nuclear foci of SUMO2/3 are a hallmark of SUMO-specific localization in mammalian cells and co-localize with a variety of nuclear proteins including the PML protein. PML nuclear bodies are implicated in nuclear stress response and reported to increase over time when cells are exposed to genotoxic stressors (Liu, Shen, Guo, Cao, & Xu, 2017). However, our short (30min) incubation with H_2_O_2_ did not statistically affect the frequency, size, or intensity of SUMO foci in either PC3 or PNT2 cells (data not shown). In summary, our data suggests that both UV irradiation and oxidative stress resulted in a rapid and concentration-dependent accumulation of SUMO in PC3 cancer cells, especially in the cytosol. These differences were not observed under similar conditions in PNT2 cells. Therefore, we focused on this novel stress-induced accumulation of SUMO conjugates specifically in the cytosol.

### Recovery from peroxide-stress was accompanied by the reduction of cytosolic SUMO levels

We evaluated the possibility that the rapid increase of cytosolic SUMO levels in PC3 cancer cells was a reversible process. PC3 and PNT2 cells were treated for 30 min with H_2_O_2_ before recovery in fresh media for 1 to 5 hours. As before, treatment with 1mM peroxide resulted in a significant and rapid increase of cytosolic (161%) and nuclear (61%) SUMO signals in PC3 cells (Fig. 4A and B). Remarkably, the nuclear SUMO signal of PC3 cells increased further and peaked 1 hour (108%) into the recovery period (Fig. 4C right panel and 4E, bottom panel). During recovery, a significant reduction of cytosolic and nuclear SUMO levels was apparent after 2 hours and SUMO levels returned to pre-treatment levels after 5 hours (Fig. 4C and 4E). Taking into account nuclear and cytosolic KmUTAG-fl signals, we found a robust RCE of SUMO that decreased steadily during recovery (Fig. 4F left panel). These data indicated a rapid and dynamic fluctuation of SUMO levels in the nucleus and cytosol of PC3 cells as they responded to and recovered from oxidative stress.

**Figure 4:**
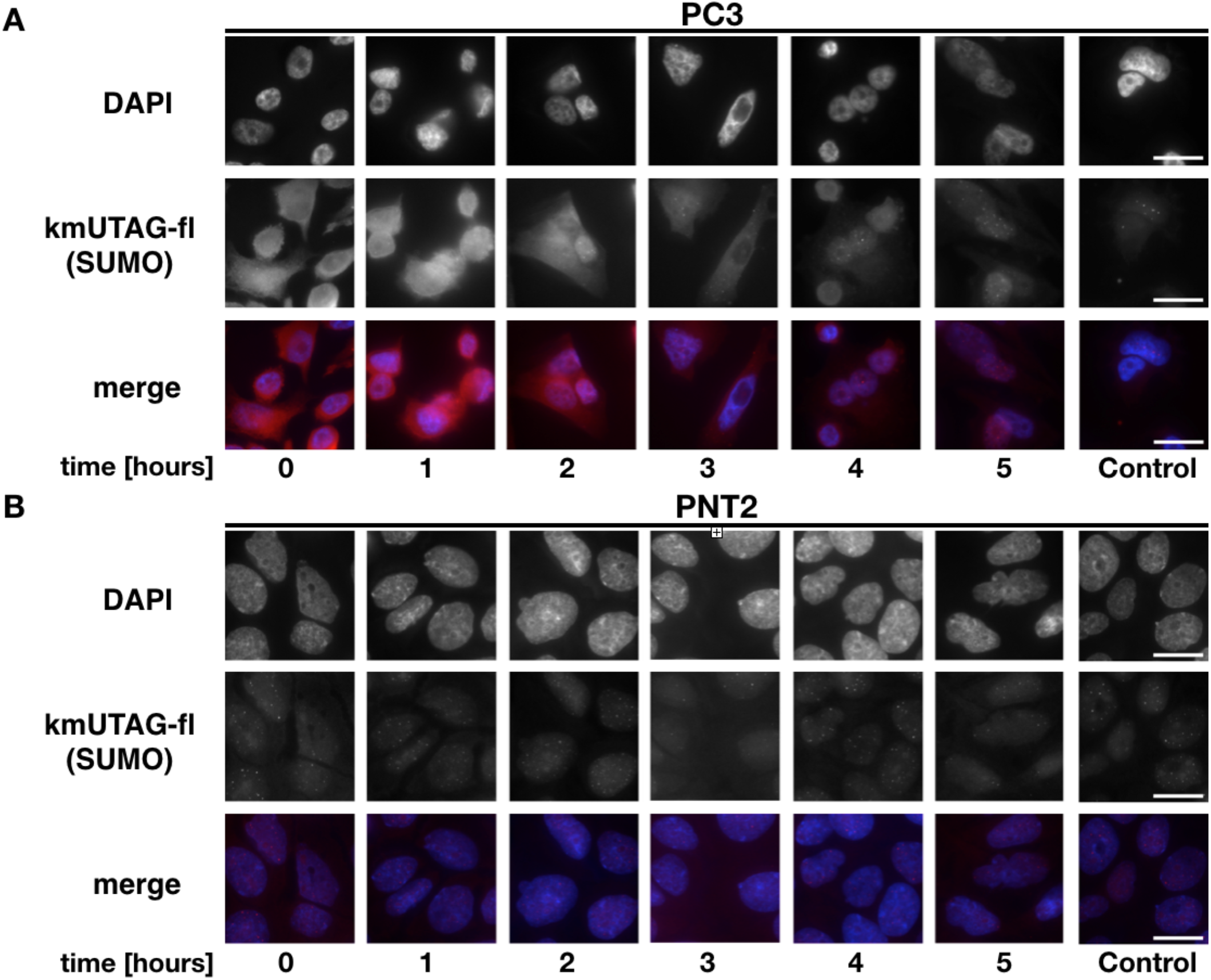
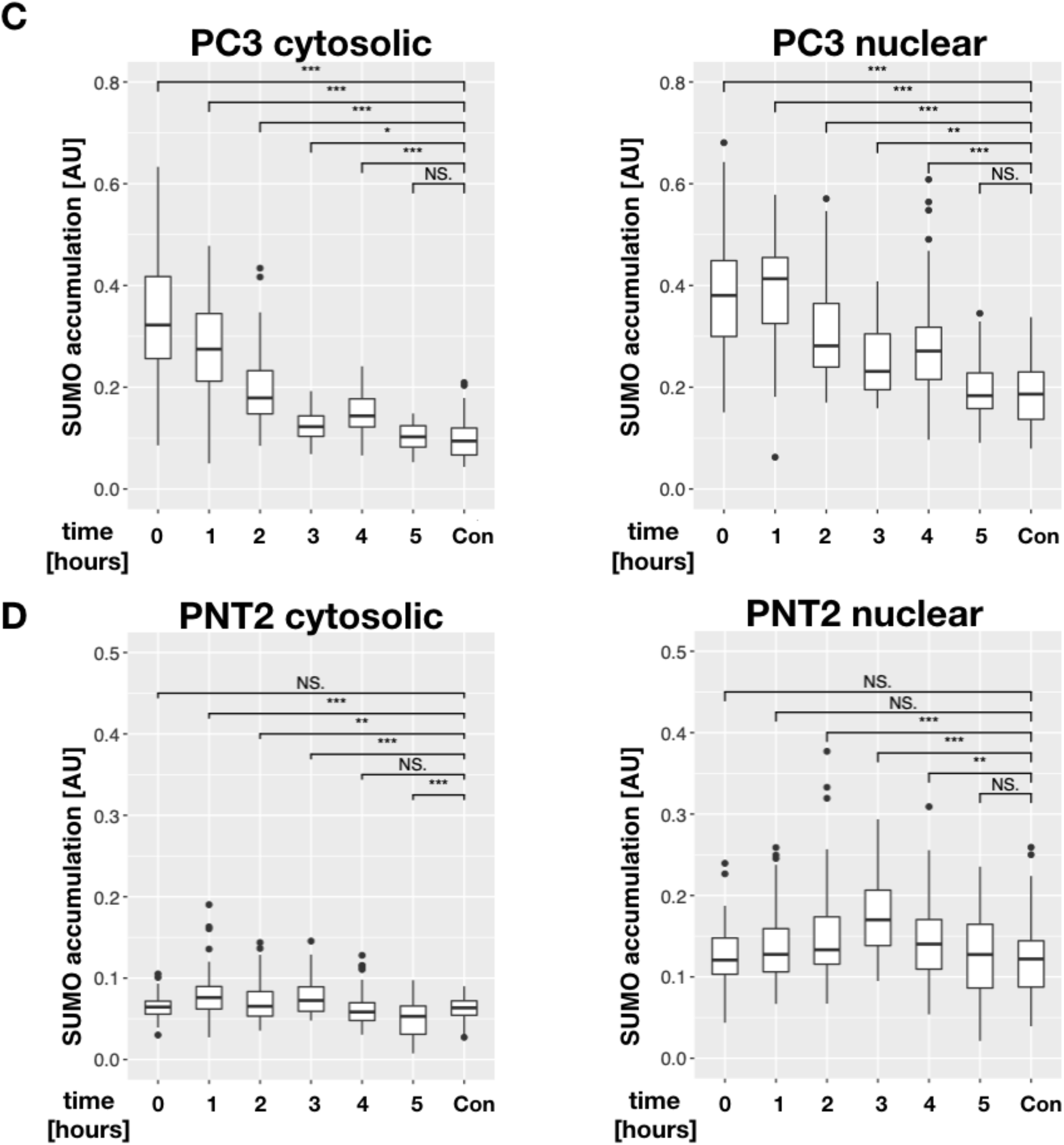
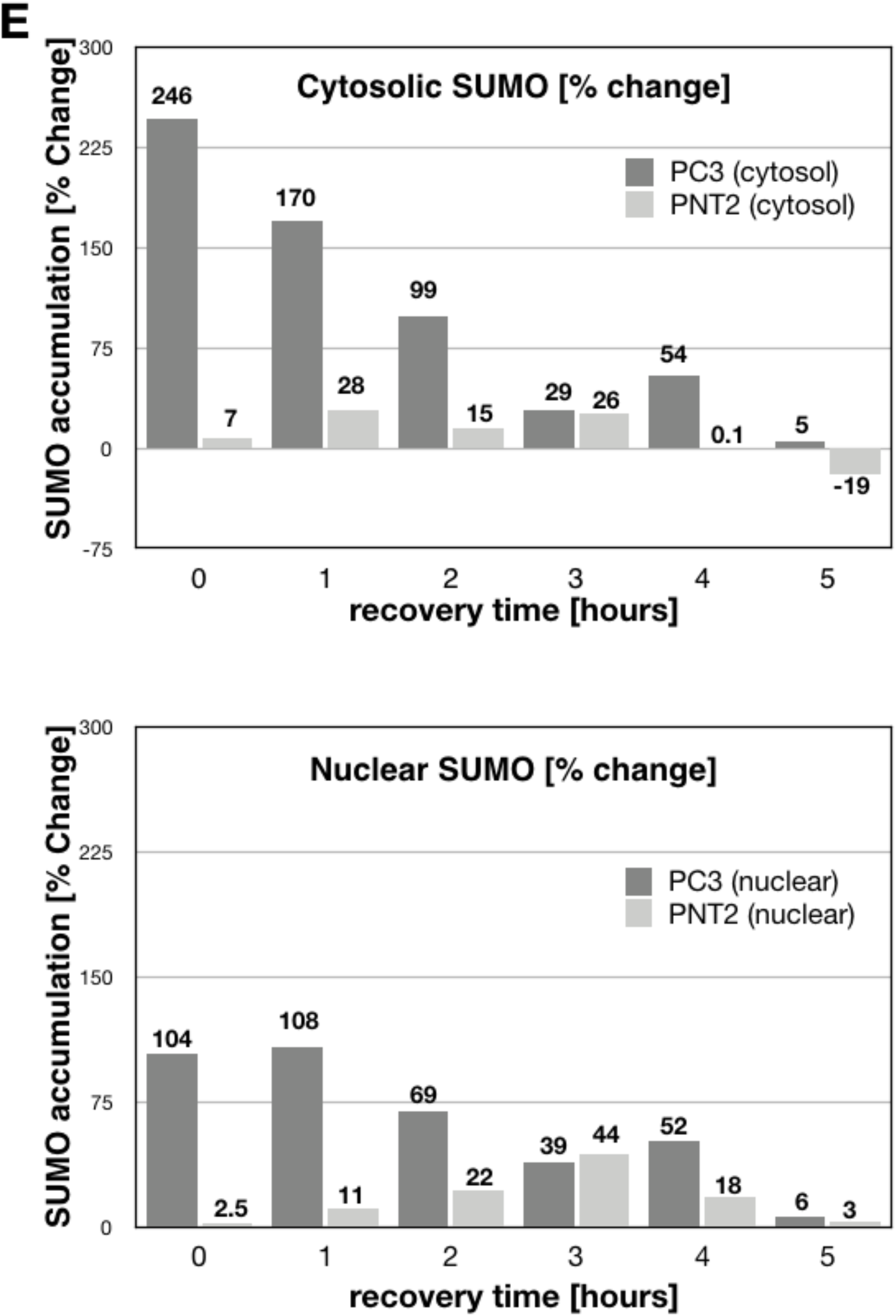
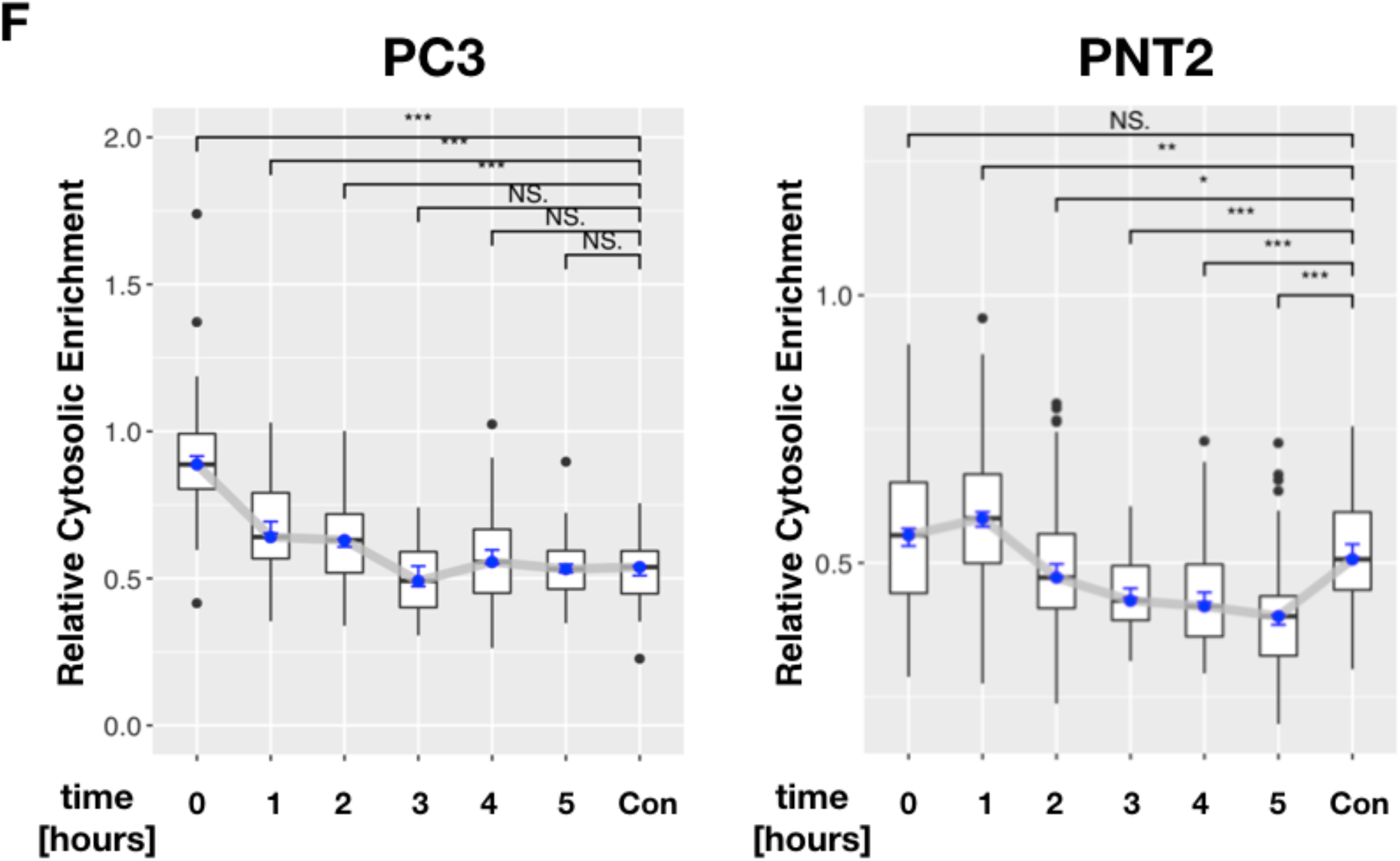
Recovery from peroxide stress is accompanied by gradually decreasing SUMO level: **(A)** H_2_O_2_ treated (1mM) or control PC3 and PNT2 cells were fixed and stained for SUMO with KmUTAG-fl after the indicated recovery times (0-5 hours). Representative cells samples are shown. Merged cells: DAPI (blue), KmUTAG-fl (red) **(B)** Same as (A) except PNT2 cells. **(C)** Decreasing KmUTAG-fl signal intensity (cytosolic and nuclear) in recovering PC3 cells (PC3 0hr n=54; PC3 1hr n=60; PC3 2hr n=93; PC3 3hr n=15; PC3 4hr n=51; PC3 5hr n=54; PC3 Control n=47). **(D)** Little or no significant change in cytosolic and nuclear KmUTAG-fl signal intensity in recovering PNT2: (0hr n=72; PNT2 1hr n=95; PNT2 2hr n=103; PNT2 3hr n=60; PNT2 4hr n=102; PNT2 5hr n=103; PNT2 Control n=70). **(E)** Comparison in the change [%] of cytosolic and nuclear (SUMO) KmUTAG-fl signal intensity in PC3 and PNT2 cells. <u>Top panel</u>: change [%] of cytosolic (SUMO) KmUTAG-fl signal. <u>Bottom panel</u>: change [%] of nuclear (SUMO) KmUTAG-fl signal. The intensity change was calculated as the intensity difference between the treated cells and control cells. **(F)** The Relative Cytosolic Enrichment (RCE) ratio of the KmUTAG-fl signal in PC3 cells (left panel) and PNT2 cells (right panel) after the indicated recovery times. Note the delayed and reduced RCE of PNT2 cells compared to PC3 cells. Statistical analysis by unpaired t-test (NS: P > 0.05, * P ≤ 0.05, ** P ≤ 0.01, *** P ≤ 0.001). Scale bars: 20 µm.

In contrast, the increase of cytosolic (28%) and nuclear (44%) SUMO levels in PNT2 cells was significantly less pronounced (compare Fig. 4D and 4E). Notably the cytosolic and the nuclear SUMO signal of PNT2 cells peaked between 1 and 3 hours into the recovery period, suggesting slower SSR response kinetics in PNT2 cells (Fig. 4D and 4E). This was also apparent when comparing the RCE of PC3 and PNT2 cells. The RCE of SUMO in PC3 cells peaked immediately after H_2_O_2_ treatment (0.85) while the RCE of SUMO in PNT2 cells peaked after 1 hour (0.58, Fig. 4F). In summary, these data suggest that SUMO levels of the PC3 prostate cancer cell line significantly increased after H_2_O_2_-induced stress and subsequently decreased as part of a recovery process.

### Stress-induced increase of cytosolic SUMO levels was observed in LNCaP prostate cancer cells with low metastatic potential

Our finding that stress rapidly increased cytosolic SUMO levels in the highly aggressive PC3 cell line prompted us to question whether this accumulation of SUMO is linked to metastatic potential. To assess this possibility, we compared stress-induced SUMO levels in the LNCaP cell line, possessing low metastatic potential, to the highly aggressive PC3 and the non-malignant PNT2 cells (Spans et al., 2014). For this experiment, we simultaneously treated LNCaP, PC3, and PNT2 cells with H_2_O_2_ (1mM, 30min) as detailed above (Fig. 3 and 4) and compared cytosolic and nuclear SUMO levels before and after treatment. SUMO levels in nuclei and cytosol of untreated LNCaP, PC3, and PNT2 cells showed different levels of SUMO before treatment (Fig 5A). After H_2_O_2_ treatment, cytosolic SUMO levels of all cell lines increased significantly (Fig. 5B left). The largest increase in cytosolic SUMO signal after peroxide exposure was detected in LNCaP cells (71%), followed by PC3 (44%), and lastly PNT2 cells (13%) (Fig 5B and C). Therefore, cytosolic SUMO levels in LNCaP cells and in PC3 cells were 5.5 and 3.5-fold higher respectively than PNT2 cells after exposure. As before, a small but significant change in nuclear SUMO levels was detected in PC3 and PNT2 cell lines: SUMO levels were decreased by 10.3% and 7.8% below the untreated control samples, respectively (Fig. 4C). Concomitantly, H_2_O_2_-treated LNCaP cells did not show a significant change in nuclear SUMO levels (Fig. 5B right). Reduced nuclear SUMO levels after peroxide exposure were likely an authentic effect, as a large number of cells (n>300 for each) were scored for this comparison. Importantly, taking into account nuclear and cytosolic KmUTAG signals, a robust increase of the RCE of SUMO was detected for both LNCaP and PC3 cells (64% increase for LNCaP and 55% increase for PC3 cells) but this change was much less pronounced in normal PNT2 cells (27% increase) (Fig. 5D). Importantly, the RCE of PC3 and LNCaP cells showed similar slopes and magnitude, suggesting the observed cytosolic enrichment of SUMO may not be related to the differences in cancerous potential. In summary, our data obtained with the KmUTAG-fl SUMO biosensor indicate that the detected increase in the RCE of SUMO levels is a hallmark of the SSR in cancerous cells.

**Figure 5:**
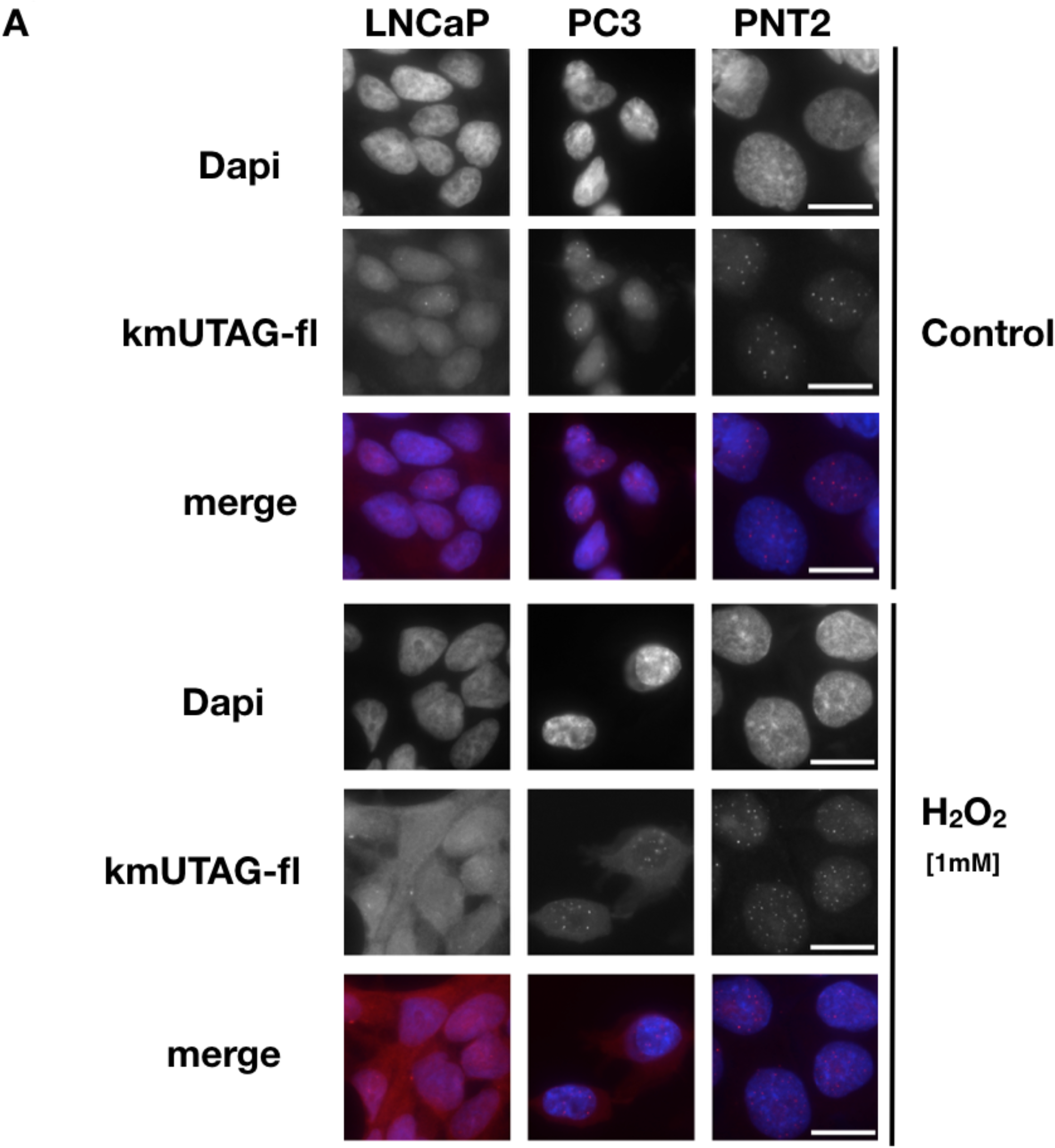
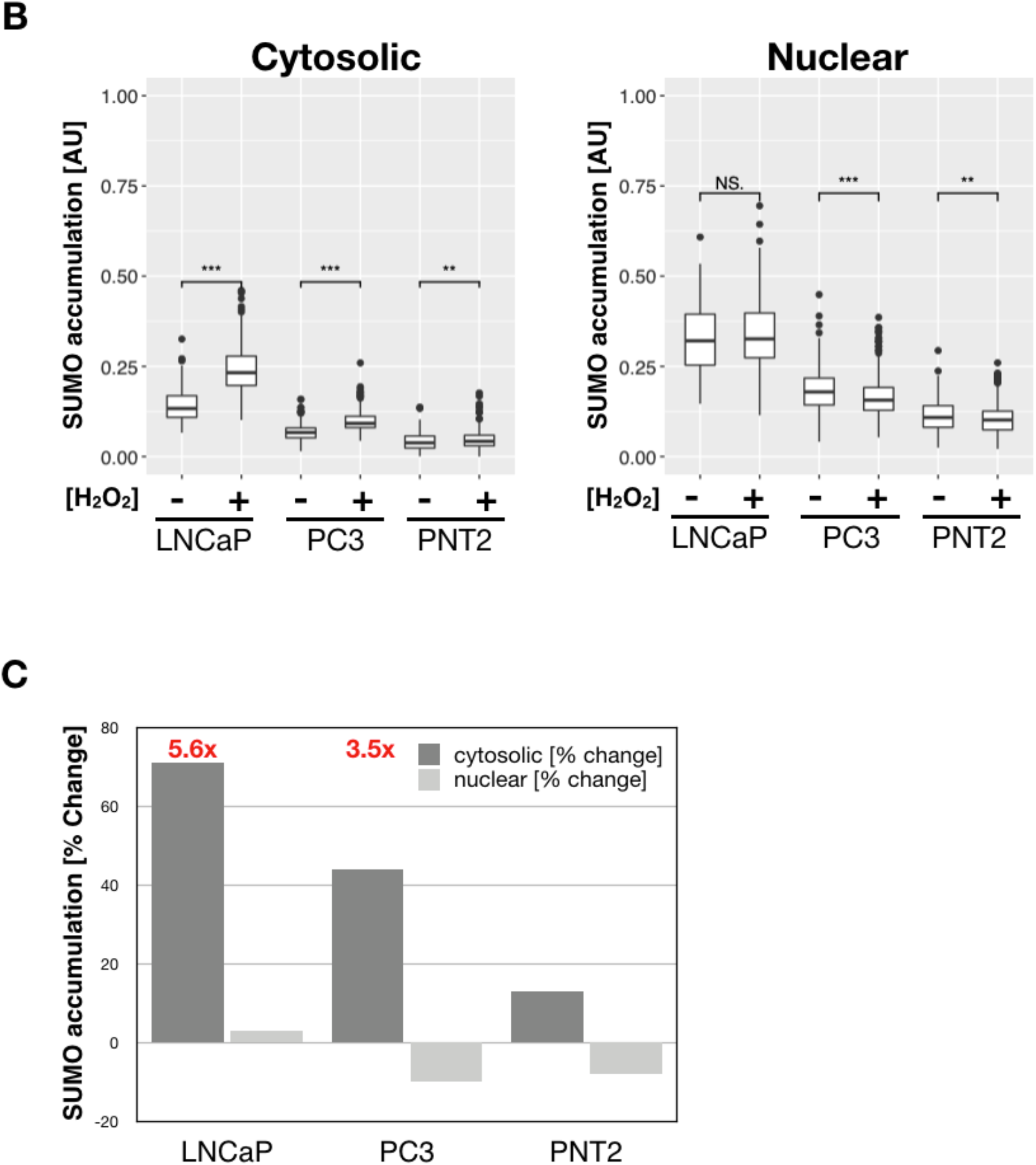
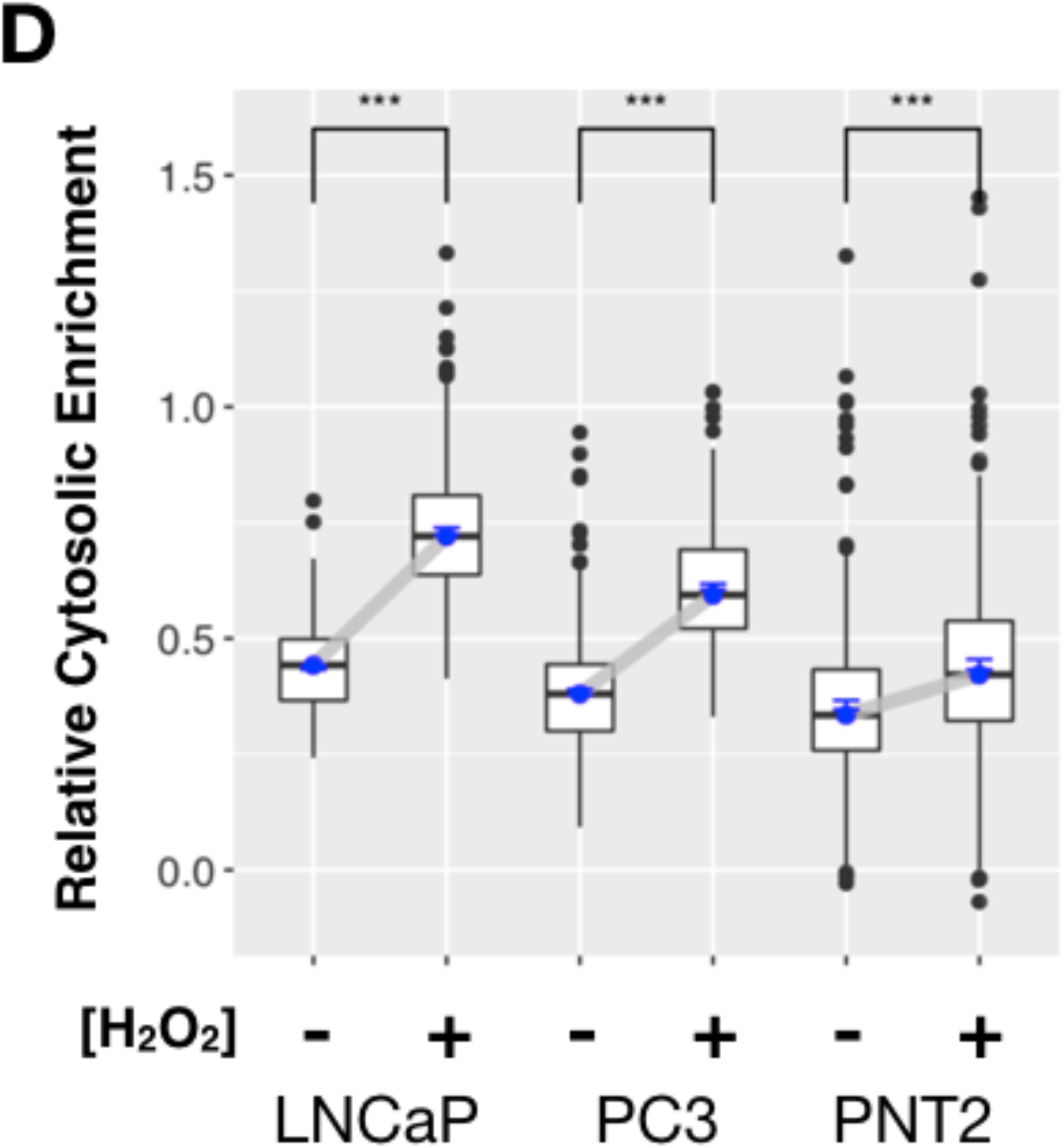
H_2_O_2_ stress induces increase in cytosolic SUMO levels in LNCaP cells with low-metastatic potential. **(A)** H_2_O_2_ treated (1mM) or Control LNCaP, PC3 and PNT2 cells were fixed and stained for SUMO with KmUTAG-fl. Representative cells of each cell line are shown. Merged cells: DAPI (blue) and KmUTAG-fl (red). **(B)** KmUTAG-fl signal intensity in nuclei and cytosol of LNCaP, PC3 and PNT2 cells across treatment groups. (LNCaP_Control n=402; LNCaP_Peroxide n=345; PC3_Control n=368; PC3_Peroxide n=422; PNT2_Control n=342; PNT2_Peroxide n=331). **(C)** The largest change [%] in KmUTAG-fl signal intensity is observed in nuclei and cytosol of LNCaP and PC3 cells when compared to PNT2 cells. The intensity change was calculated as the intensity difference between the peroxide-treated cells and control cells. **(D)** The relative cytosolic enrichment (RCE) ratio of the KmUTAG-fl signal is most pronounced in LNCaP, PC3 when compared to PNT2 cells. Statistical analysis by unpaired t-test (NS: P > 0.05, * P ≤ 0.05, ** P ≤ 0.01, *** P≤ 0.001). Scale bars: 20 µm.

## DISCUSSION

The KmUTAG-fl biosensor reports on the presence and distribution of untagged, native SUMO conjugates in a variety of eukaryotic cells (Yin et al., 2019). Here, we used KmUTAG-fl to detect, quantitate, and analyze the cellular distribution of SUMO conjugates before and after exposure to acute proteotoxic and genotoxic stress. Using this approach, we detected significant, stress-induced differences in the distribution of cytosolic (extra-nuclear) SUMO between a normal (PNT2) and two cancer cell lines (PC3 and LNCaP). While the increase in cellular SUMO conjugate levels in response to stress has previously been observed, it is unknown how the SSR propagates throughout the cell or the extent of extra-nuclear SUMO distribution changes when cells undergo acute stress. In this report, we observed a significant increase in cytosolic SUMO in response to oxidative and UV-irradiation stress within 30 min of exposure, which was dependent on the concentration of the stressor and the time after stress exposure. Additionally, after recovery cytosolic SUMO levels returned to normal levels within a 5-hour interval. Nuclear SUMO levels were also altered during the SSR but overall the amplitude of this stress-induced modulation was lower or not significantly changed in the cell lines tested. The increase of cytosolic SUMO in response to peroxide treatment ranged from 12.6% to 70.6% (average increase ∼5-fold) and was statistically significant for both PC3 and LNCaP cell lines when compared to PNT2 cells (unpaired *t*-test), indicating this stress-induced effect was consistent and reproducible. Overall, our findings demonstrate that the SUMO biosensor KmUTAG-fl can differentiate between the SSR of cancerous and normal cells. Additionally, our work offers new insights into the dynamics of the SSR and how cancer cells modulate SUMO levels in response to stress. Ultimately, we hope to determine whether this enhanced SSR response facilitates the robustness of cancer cells.

Cancer cells are known to modulate their SUMO dynamics as part of a strategy to become more stress tolerant (Seeler & Dejean, 2017). Cancer cells are under constant threat of adverse conditions, including hypoxia within tumors, immune invasions, and aneuploidies that threaten protein homeostasis (Muz, la Puente, Azab, & Azab, 2015; Oromendia & Amon, 2014). Thus, cancer cells require enhanced stress response pathways to mitigate these effects and to maintain proteostasis and genome integrity. For our study, we used three different cell lines to observe the SSR and to identify unique cell-type specific features. PNT2 cells were established by SV40-mediated immortalization of normal adult prostatic epithelial cells, PC3 cells represent an adenocarcinoma cell line with high metastatic potential, and the LNCaP cell line has low metastatic potential. These human-derived prostate cell lines differ not only in their tumorigenicity but also their chromosomal make-up. PNT2 cells are non-malignant normal prostate epithelium immortalized with SV40, PC3 have 62 chromosomes (Tai et al., 2011), and LNCaP harbor 79-91 chromosomes (Horoszewicz et al., 1983). Therefore, both the expression levels and copy number of SUMO genes are increased in these tumorigenic cell lines (Kerscher unpublished results). Our observation that cytosolic SUMO levels are significantly increased after stress exposure of PC3 and LNCaP cells may be linked to the increased expression levels of SUMO in comparison to PNT2 cells. However, it is important to note that de-novo protein synthesis (translation) is not required for the SSR to proceed (Lewicki et al., 2014). This implies that SUMO levels *per se* do not increase in stress-exposed cells. Rather, pre-existing SUMO is dynamically transferred to specific proteins in response to stress exposure and as part of transcriptional re-programming in the nucleus. Therefore, it is not surprising that overall SUMO levels do not increase in the nuclei of stressed cells and our data are thus consistent with previous SSR studies.

Regarding the observed stress-dependent cytosolic enrichment of SUMO, one possibility is that after stress sumoylated nuclear proteins may “spill” into the cytosol. This is unlikely because jn our study SUMO levels do not simply equilibrate across the cell (e.g. see Fig 3E). We posit that the apparent stress-dependent cytosolic enrichment of SUMO is most likely due to conversion of free (or inaccessible) SUMO into readily detectable SUMO conjugates and chains. Both the mechanism and relevant targets of the SSR in the cytosol have not been worked out at this point. Work in yeast has established that Siz/PIAS SUMO E3 ligases drive the SSR and SUMO protease Ulp2 is required for inactivation of the SSR (Takahashi *et al*., 2008, Lewicki *et al*., 2015). Some SUMO E3 ligases (e.g. PIAS4) and SUMO proteases (e.g. SENP2) localize to the nucleus as well as the cytosol, suggesting that the observed increase and decrease of SUMO conjugates may take place outside of the nucleus without transport across the nuclear envelope.

Next to SUMO, SUMO E1, E2, and E3 levels are also altered in cancer cells and this likely affects the dynamics of stress-induced sumoylation. For example, the SUMO E2 ligase Ubc9 is upregulated in breast, lung, neck and head cancer specimens, and is elevated 5.7-fold in breast tumors compared to normal breast tissues (Wu *et al*., 2009). Ubc9 may represent a useful biomarker for cervical cancer and is linked to the progression of HPV’s oncogenicity (Mattoscio *et al*., 2015). Hundreds of proteins in the cytosol may be targets of stress-induced sumoylation. As outlined below, sumoylation may serve to stabilize proteins during acute proteotoxic stress. Additionally, chains of SUMO monomers likely make up a considerable part of stress-induced sumoylated products (Srikumar et al., 2013). The identification of these proteins, while of interest, is beyond the scope of the present analysis.

Sumoylation is considered a predominantly nuclear event, so the rapid stress-induced increase and decrease of SUMO conjugates in the cytosol of cancer cells is an interesting finding. A recent review on sub-cellular sumoylation in the heart posits that sumoylation in the extra-nuclear compartment of cardiomyocytes is generally cardio-protective (Le, Martin, Fujiwara, & Abe, 2017). More specifically, results from a study on SUMO2/3 suggest that stress-induced sumoylation serves to temporarily stabilize (keep soluble) misfolded proteins and targets those that can’t be refolded for proteasomal degradation (Liebelt et al., 2019). Nevertheless, most information on stress-induced sumoylation concerns its nuclear effects and ignores the cytoplasmic roles of SUMO. For example, SUMO isoforms may serve a chaperone-like function to maintain the homeostasis of large chromatin-associated nuclear proteins during stress (Seifert et al., 2015) and stress-induced sumoylation is required for transcriptional re-programming (Lewicki et al., 2015). In this respect, our observation of a transient increase of SUMO due to acute UV and oxidative stress underscores the dynamic nature of the SSR in the cell. Additionally, it is an important reminder that the effects of SUMO (and especially the SSR in normal and transformed cell lines) are likely to produce very different, cell line-specific outcomes. Many proteomics studies on the SSR in mammalian cell lines are conducted in cancer-derived cell lines and these studies do not always include normal, immortalized comparators.

In summary, using the pan-SUMO specific KmUTAG-fl biosensor, we have identified a transient, reversible, stress-induced increase of SUMO conjugates in the cytosol of PC3 and LNCaP cells. This stress-induced cytosolic SUMO enrichment clearly distinguishes PC3 and LNCaP cells from normal immortalized PNT2 cells, suggesting that it may be part of a stress-tolerance pathway that is specific for cancer cells. One of our future goals to identify these stress-induced SUMO targets and conjugates after cell fractionation. The latter is technically difficult as SUMO is rapidly deconjugated or degraded during cell fractionation.

While the cytosolic increase of SUMO in response to acute stress may be a feature of these prostate cancer cells, we have not yet shown that this enhanced SSR is correlated with enhanced oncogenic potential (defined as an increase in cellular migration, invasiveness, and aggressive cell proliferation). Looking forward, we will investigate a potential link between transiently increased SUMO levels and oncogenic potential that would underscore the importance that the SSR plays during cellular transformation. Additionally, to further understand how the SSR unfolds in cancer cells, we plan to use a cell-penetrating CPP-adaptor system to deliver and release the KmUTAG-fl biosensor into the cytoplasm of mammalian cells (Salerno et al., 2016). The successful delivery of KmUTAG-fl will allow us to visualize the propagation of the SSR inside living cells and study the SUMO-specific features as cells go from non-cancer to malignant. These experiments chart an important new route in our studies of the SSR and ultimately will help us understand the role that SUMO plays in the enhanced stress tolerance of eukaryotic pathogens and abnormal cells.

## ACKNOWLEDGEMENTS

We would like to thank the present and past members of the Kerscher lab for their support especially Jen Peek, and Shriie Ganesh. Additionally, we would like to thank members of the William & Mary Biology Department for their support, intellectual input, critical reading of the manuscript, or technical support, specifically Diane Shakes, Shanta Hinton, and Lidia Epp. This work was supported by the Commonwealth Research Commercialization Fund (CRCF) award to OK, the Bailey-Huston Research Fund, and the HHMI Science and Research Program at William and Mary to RY.

## MATERIALS & METHODS

### Cell Culture and Maintenance

PC3, PNT2, and LNCaP cells were grown in RPMI media with 10% heat inactivated FBS (Thermo Fisher Scientific #10438018) and 1% antifungal/antibiotic (anti/anti) (Thermo Fisher Scientific #15240062). All cells were grown at 37°C in a humidified incubator which is kept constant at 5% CO_2_.

### Expression of KmUTAG-fl and in vitro Assays

A codon-optimized bacterial overexpression clone of mCherry-KmUTAG was generated as previously described (Yin et al., 2019). KmUTAG-fl biosensor were over-expressed in BL21-STAR(DE3) cells (Muench *et al*., 2003). Purification of KmUTAG-fl in these cells was as previously described (Yin et al., 2019) except that KmUTAG-fl was bound to magnetic SPOT-trap beads (Chromotek). To assess SUMO-trapping activity of KmUTAG-fl, SUMO-binding reactions were performed on SUMO1 beads and with recombinant SUMO-CAT fusion protein (Peek et al., 2018).

For SUMO staining, PC3, PNT2 or LNCaP cells were grown to 80% confluency and counted using a hemocytometer. Each well in a 6-well plate (Fisher Scientific 07-200-83) received 300,000 cells in 2 mL media. Cells were incubated for 24 hours until 80% confluent. After fixing the cells with fresh 4% paraformaldehyde (PF) for 20 minutes at room temperature, cells were permeabilized for 15 min with 0.1% Triton X-100 in dPBS. Next, the cells were incubated with 0.1M glycine-HCL (pH 2.0) for 10 seconds. The pH was immediately neutralized with 500μL of 10X SPB (500mM Tris-HCL, pH 8.0 2% NP-40, 1.5M NaCl) (Peek et al., 2018) and coverslips were removed from the well and placed into humidity chambers. For KmUTAG-fl staining, 2 ug of recombinant KmUTAG-fl was added to 100μL 1X SPB containing 5mM TCEP. The mix was transferred onto the coverslip and incubated at room temperature for 1 hr in the humidity chamber. For KmUTAG-fl and anti-SUMO2/3 antibody co-staining, 2 ug of KmUTAG-fl and 0.5 μL SUMO2/3 8A2 (obtained for Developmental Studies Hybridoma Bank - DSHB Hybridoma Product SUMO-2 8A2 (X.-D. Zhang et al., 2008)) were mixed with 100 μL blocking buffer, pipetted onto the coverslip, and incubated in room temperature for 1 hr. After washing the coverslips with dPBS, 0.5 μL anti-mouse Alexa Fluor 488–conjugated antibody (Jackson ImmunoResearch 115-545-003) in 100μL blocking buffer (Prometheus 20-313), was pipetted onto the coverslip of KmUTAG-fl and SUMO2/3 co-staining slides, and incubated at room temperature for 1 hr. After washes with dPBS, the coverslips were inverted onto pre-cleaned microscopy slides with FLUORO-GEL II with DAPI (Electron Microscopy Sciences 50-246-93). The slides were stored at -20°C overnight before viewing under the microscope.

### Stress Treatment

To test the SUMO stress sesponse (SSR) to UV damage, cells were grown until ∼80% confluent on coverslips and subjected to UV irradiation before proceeding with fixation and staining. The coverslips were first transferred from the 6-well plate to a humidity chamber. Excess media on the coverslip was removed and the humidity chamber containing the coverslip was put into a UV chamber (GS Gene Linker). The UV intensity was adjusted per manufacturer’s protocol. After irradiation, coverslips were immediately placed back into culture medium and placed back into the tissue culture incubator for an additional 30 min. Fixation and staining was completed as detailed above.

For peroxide stress treatment, H_2_O_2_ was added from a 3M stock solution to 2 mL cultures in a 6-well plate to achieve the desired final concentration. Cells were then incubated for an additional 30 min in the tissue culture incubator. After incubation, the tissue culture supernatant was removed and the cells were washed in dPBS before fixation and staining with KmUTAG-fl. All experiments were completed at least in duplicate and include 2-10 slides with cells per condition tested. Representative images are shown after normalization.

### Microscopy and Data Analysis

Images were acquired using a fully automated Nikon A1R inverted confocal microscope or a Zeiss Axioscope using the appropriate filter sets. Fluorescence staining intensity was quantified using CellProfiler (www.cellprofiler.org (McQuin et al., 2018)). Nuclei and cytosol of imaged cells were automatically detected as DAPI-stained nuclei and kmUTAG-fl stained nuclei and cytosol, respectively. For normalization between images, the background fluorescence intensity between cell features was subtracted from the mean intensity of each compartment of individual cells to obtain the cytoplasmic and nuclear staining intensity. The relative cytosolic enrichment was calculated as the ratio between cytoplasmic and nuclear staining intensities per cell. Number of cells used for evaluation is listed in the individual figure legends. Unpaired parametric t-test was used to assess significant differences in staining intensity of cytosolic and nuclear kmUTAG-fl before and after stress exposure. Significance level signs were displayed to indicate the result of the T-test (NS: P > 0.05, * P ≤ 0.05, ** P ≤ 0.01, *** P ≤ 0.001). The percentage difference between mean fluorescence intensity levels per treatment group was calculated to further describe the change in kmUTAG-fl signal intensity before and after stress. Data was graphed using R software.

## References

Aurich-Costa, J., Vannier, A., Grégoire, E., Nowak, F., & Cherif, D. (2001). IPM-FISH, a new M-FISH approach using IRS-PCR painting probes: application to the analysis of seven human prostate cell lines. Genes, Chromosomes & Cancer, 30(2), 143– 160.

Berthon, P., Cussenot, O., Hopwood, L., Leduc, A., & Maitland, N. (1995). Functional expression of sv40 in normal human prostatic epithelial and fibroblastic cells - differentiation pattern of nontumorigenic cell-lines. International Journal of Oncology, 6(2), 333–343. http://doi.org/10.3892/ijo.6.2.333

Bossis, G., & Melchior, F. (2006). Regulation of SUMOylation by Reversible Oxidation of SUMO Conjugating Enzymes. Molecular Cell, 21(3), 349–357. http://doi.org/10.1016/j.molcel.2005.12.019

Golebiowski, F., Matic, I., Tatham, M. H., Cole, C., Yin, Y., Nakamura, A., et al. (2009). System-wide changes to SUMO modifications in response to heat shock. Science Signaling, 2(72), ra24–ra24. http://doi.org/10.1126/scisignal.2000282

Hendriks, I. A., D’Souza, R. C. J., Yang, B., Verlaan-de Vries, M., Mann, M., & Vertegaal, A. C. O. (2014). Uncovering global SUMOylation signaling networks in a site-specific manner. Nature Structural & Molecular Biology, 21(10), 927–936. http://doi.org/10.1038/nsmb.2890

Horoszewicz, J. S., Leong, S. S., Kawinski, E., Karr, J. P., Rosenthal, H., Chu, T. M., et al. (1983). LNCaP model of human prostatic carcinoma. Cancer Research, 43(4), 1809–1818.

Karami, S., Lin, F.-M., Kumar, S., Bahnassy, S., Thangavel, H., Quttina, M., et al. (2017). Novel SUMO-Protease SENP7S Regulates β-catenin Signaling and Mammary Epithelial Cell Transformation. Scientific Reports, 7(1), 46477. http://doi.org/10.1038/srep46477

Kerscher, O., Felberbaum, R., & Hochstrasser, M. (2006). Modification of proteins by ubiquitin and ubiquitin-like proteins. Cell and Developmental Biology, 22, 159– 180. http://doi.org/10.1146/annurev.cellbio.22.010605.093503

Knipscheer, P., van Dijk, W. J., Olsen, J. V., Mann, M., & Sixma, T. K. (2007). Noncovalent interaction between Ubc9 and SUMO promotes SUMO chain formation. EMBO Journal, 26(11), 2797–2807. http://doi.org/10.1038/sj.emboj.7601711

Le, N.-T., Martin, J. F., Fujiwara, K., & Abe, J.-I. (2017). Sub-cellular localization specific SUMOylation in the heart. Biochimica Et Biophysica Acta. Molecular Basis of Disease, 1863(8), 2041–2055. http://doi.org/10.1016/j.bbadis.2017.01.018

Lewicki, M. C., Srikumar, T., Johnson, E., & Raught, B. (2015). The S. cerevisiae SUMO stress response is a conjugation-deconjugation cycle that targets the transcription machinery. Journal of Proteomics, 118, 39–48. http://doi.org/10.1016/j.jprot.2014.11.012

Liebelt, F., Sebastian, R. M., Moore, C. L., Mulder, M. P. C., Ovaa, H., Shoulders, M. D., & Vertegaal, A. C. O. (2019). SUMOylation and the HSF1-Regulated Chaperone Network Converge to Promote Proteostasis in Response to Heat Shock. Cell Reports, 26(1), 236–249.e4. http://doi.org/10.1016/j.celrep.2018.12.027

Liu, S.-B., Shen, Z.-F., Guo, Y.-J., Cao, L.-X., & Xu, Y. (2017). PML silencing inhibits cell proliferation and induces DNA damage in cultured ovarian cancer cells. Biomedical Reports, 7(1), 29–35. http://doi.org/10.3892/br.2017.919

Mattoscio, D. (2015). The SUMO conjugating enzyme UBC9 as a biomarker for cervical HPV infections. Ecancermedicalscience, 9. http://doi.org/10.3332/ecancer.2015.534

McQuin, C., Goodman, A., Chernyshev, V., Kamentsky, L., Cimini, B. A., Karhohs, K. W., et al. (2018). CellProfiler 3.0: Next-generation image processing for biology. PLoS Biology, 16(7), e2005970. http://doi.org/10.1371/journal.pbio.2005970

Muz, B., la Puente, de, P., Azab, F., & Azab, A. K. (2015). The role of hypoxia in cancer progression, angiogenesis, metastasis, and resistance to therapy. Hypoxia (Auckland, N.Z.), 3, 83–92. http://doi.org/10.2147/HP.S93413

Oromendia, A. B., & Amon, A. (2014). Aneuploidy: implications for protein homeostasis and disease. Disease Models & Mechanisms, 7(1), 15–20. http://doi.org/10.1242/dmm.013391

Peek, J., Harvey, C., Gray, D., Rosenberg, D., Kolla, L., Levy-Myers, R., et al. (2018). SUMO targeting of a stress-tolerant Ulp1 SUMO protease. PLoS ONE, 13(1), e0191391. http://doi.org/10.1371/journal.pone.0191391

Pinto, M. P., Carvalho, A. F., Grou, C. P., Rodríguez-Borges, J. E., Sá-Miranda, C., & Azevedo, J. E. (2012). Heat shock induces a massive but differential inactivation of SUMO-specific proteases. Biochimica Et Biophysica Acta, 1823(10), 1958– 1966. http://doi.org/10.1016/j.bbamcr.2012.07.010

Saitoh, H., & Hinchey, J. (2000). Functional heterogeneity of small ubiquitin-related protein modifiers SUMO-1 versus SUMO-2/3. The Journal of Biological Chemistry, 275(9), 6252–6258.

Salerno, J. C., Ngwa, V. M., Nowak, S. J., Chrestensen, C. A., Healey, A. N., & McMurry, J.L. (2016). Novel cell penetrating peptide-adaptors effect intracellular delivery and endosomal escape of protein cargos. Journal of Cell Science. http://doi.org/10.1242/jcs.182113

Sarge, K. D., & Park-Sarge, O.-K. (2009). Sumoylation and human disease pathogenesis. Trends in Biochemical Sciences, 34(4), 200–205. http://doi.org/10.1016/j.tibs.2009.01.004

Seeler, J.-S., & Dejean, A. (2017). SUMO and the robustness of cancer. Nature Reviews Cancer, 17(3), 184–197. http://doi.org/10.1038/nrc.2016.143

Seifert, A., Schofield, P., Barton, G. J., & Hay, R. T. (2015). Proteotoxic stress reprograms the chromatin landscape of SUMO modification. Science Signaling, 8(384), rs7–rs7. http://doi.org/10.1126/scisignal.aaa2213

Spans, L., Helsen, C., Clinckemalie, L., Van den Broeck, T., Prekovic, S., Joniau, S., et al. (2014). Comparative genomic and transcriptomic analyses of LNCaP and C4-2B prostate cancer cell lines. PLoS ONE, 9(2), e90002. http://doi.org/10.1371/journal.pone.0090002

Srikumar, T., Lewicki, M. C., Costanzo, M., Tkach, J. M., van Bakel, H., Tsui, K., et al. (2013). Global analysis of SUMO chain function reveals multiple roles in chromatin regulation. The Journal of Cell Biology, 201(1), 145–163. http://doi.org/10.1083/jcb.201210019.dv

Tai, S., Sun, Y., Squires, J. M., Zhang, H., Oh, W. K., Liang, C.-Z., & Huang, J. (2011). PC3 is a cell line characteristic of prostatic small cell carcinoma. The Prostate, 71(15), 1668–1679. http://doi.org/10.1002/pros.21383

Wang, Q., Xia, N., Li, T., Xu, Y., Zou, Y., Zuo, Y., et al. (2013a). SUMO-specific protease 1 promotes prostate cancer progression and metastasis. Oncogene, 32(19), 2493– 2498. http://doi.org/10.1038/onc.2012.250

Wang, R.-T., Zhi, X.-Y., Zhang, Y., & Zhang, J. (2013b). Inhibition of SENP1 induces radiosensitization in lung cancer cells. Experimental and Therapeutic Medicine, 6(4), 1054–1058. http://doi.org/10.3892/etm.2013.1259

Wu, F., Zhu, S., Ding, Y., Beck, W. T., & Mo, Y.-Y. (2009). MicroRNA-mediated regulation of Ubc9 expression in cancer cells. Clinical Cancer Research, 15(5), 1550–1557. http://doi.org/10.1158/1078-0432.CCR-08-0820

Yin, R., Harvey, C., Shakes, D. C., & Kerscher, O. (2019). Localization of SUMO-modified Proteins Using Fluorescent Sumo-trapping Proteins. Journal of Visualized Experiments: JoVE, (146), e59576. http://doi.org/10.3791/59576

Zhang, X.-D., Goeres, J., Zhang, H., Yen, T. J., Porter, A. C. G., & Matunis, M. J. (2008). SUMO-2/3 modification and binding regulate the association of CENP-E with kinetochores and progression through mitosis. Molecular Cell, 29(6), 729–741. http://doi.org/10.1016/j.molcel.2008.01.013

